# Genome-wide CRISPR/Cas9 screen reveals factors that influence the susceptibility of tumor cells to NK cell-mediated killing

**DOI:** 10.1101/2024.10.08.615667

**Authors:** Sophie Guia, Aurore Fenis, Justine Galluso, Hakim Medjouel, Bertrand Escalière, Angelica Modelska, Margaux Vienne, Noella Lopes, Amélie Pouchin, Benjamin Rossi, Laurent Gauthier, Sandrine Roulland, Eric Vivier, Emilie Narni-Mancinelli

## Abstract

**Background:** Natural killer (NK) cells exhibit potent cytotoxic activity against various cancer cell types. Over the past five decades, numerous methodologies have been employed to elucidate the intricate molecular mechanisms underlying NK cell-mediated tumor control. While significant progress has been made in elucidating the interactions between NK cells and tumor cells, the regulatory factors governing NK cell-mediated tumor cell destruction are not yet fully understood. This includes the diverse array of tumor ligands recognized by NK cells and the mechanisms that NK cells employ to eliminate tumor cells.

**Methods:** In this study, we employed a genome-wide CRISPR/Cas9 screening approach in conjunction with functional cytotoxicity assays to delineate the proteins modulating the susceptibility of colon adenocarcinoma HCT-116 cells to NK cell-mediated cytotoxicity.

**Results:** Analysis of guide RNA (gRNA) distribution in HCT-116 cells that survived co-incubation with NK cells identified ICAM-1 as a pivotal player in the NKp44-mediated immune synapse, with NKp44 serving as an activating receptor crucial for the elimination of HCT-116 tumor cells by NK cells. Furthermore, disruption of genes involved in the apoptosis or IFN-γ signaling pathways conferred resistance to NK cell attack. We further dissected that NK cell-derived IFN-γ promotes mitochondrial apoptosis *in vitro* and exerts control over B16-F10 lung metastases *in vivo*.

**Conclusion:** Monitoring ICAM-1 levels on the surface of tumor cells or modulating its expression should be considered in the context of NK cell-based therapy. Additionally, considering the diffusion properties of IFN-γ, our findings highlight the potential of leveraging NK cell-derived IFN-γ to enhance direct tumor cell killing and facilitate bystander effects via cytokine diffusion, warranting further investigation.

**WHAT IS ALREADY KNOWN ON THIS TOPIC:** NK cells play a crucial role in identifying and eliminating various cancer cell types. However, the mechanisms that regulate NK cell-mediated destruction of tumor cells are not yet fully understood. This involves the array of tumor ligands that NK cells recognize and the processes they utilize to carry out tumor cell elimination.

**WHAT THIS STUDY ADDS:** Our research emphasizes the critical role of ICAM-1 in NKp44-mediated destruction of HCT-116 tumor cells. Additionally, we found that interfering with genes related to apoptosis or IFN-γ signaling pathways increased resistance to NK cell attack. We showed that IFN-γ produced by NK cells induces mitochondrial apoptosis *in vitro* and helps regulate B16-F10 lung metastases *in vivo*.

**HOW THIS STUDY MIGHT AFFECT RESEARCH, PRACTICE OR POLICY:** Given the ability of IFN-γ to diffuse, our findings suggest that NK cell-derived IFN-γ can be harnessed to directly kill tumor cells and trigger bystander effects through cytokine spread. This approach holds promise for further exploration. Additionally, assessing or manipulating ICAM-1 levels on tumor cell surfaces could enhance the effectiveness of NK cell-based therapies.

## INTRODUCTION

Natural killer (NK) cells are innate lymphoid cells known for their cytolytic activity and their ability to secrete cytokines and chemokines [1]. NK cells form immunological synapses with the target cells, leading to their destruction through the degranulation of cytotoxic molecules, a mechanism akin to that of CD8^+^ T cells. NK cell-mediated tumor cell killing also involves the expression of ligands for death domain receptors, such as FasL in humans and mice and TRAIL in humans [1, 2]. NK cells can also mediate antibody-dependent cell cytotoxicity (ADCC). In addition, NK cells secrete cytokines such as IFN-γ and TNF-α, growth factors and chemokines that influence both innate and adaptive immune responses. For instance, IFN-γ production impacts T cell responses and activates myeloid cells. Conversely, NK cells can suppress T cell responses by directly killing activated T cells or antigen-presenting cells [3].

Ample evidence highlights the role of NK cells in anti-tumor immunity through their cytotoxic activities. *In vivo* studies with perforin-deficient mice demonstrate a significant defect in NK cell ability to kill their targets [4]. *In vitro*, perforin-deficient NK cells can still kill the RMA-S (MHC-I) tumor cells but required an extended 18-hour chromium release assay, compared to the conventional 4-hour assay [5, 6]. In these conditions, the addition of an anti-FasL antibody abrogates tumor cell lysis, underscoring the importance of FasL in NK cell-mediated cytotoxicity. Several studies have explored the relationship between NK cell cytotoxic activity in the blood and cancer risk in humans [7, 8]. A correlation between low NK cell-mediated cytotoxicity and the incidence of breast cancer suggests that defects in NK cell function may contribute to the development of breast tumors [7]. Furthermore, an 11-year follow-up study in the Japanese general population revealed that individuals with low NK cell cytotoxic activity in peripheral blood had a higher risk of cancer [9]. Additional research indicates that the number of NK cells and their ability to produce granzyme B are inversely related to disease severity in patients with chronic lymphocytic leukemia (CLL) and diffuse large B-cell lymphoma [10, 11], and positively associated with a good prognosis in Hodgkin’s lymphoma [12].

NK cells sense their environment and recognize tumor cells by integrating signals from ligands that bind to inhibitory and activating germline-encoded receptors [1]. Activation of NK cells results from engagement of activating receptors, including natural cytotoxicity receptors (NCRs) such as NKp46 in human and mice, and NKp30 and NKp44 in humans, signaling lymphocyte-activating molecule (SLAM)-related receptors and NKG2D [13]. These interactions initiate various signaling pathways that lead to NK cell activation. The NCRs recognize bacterial and tumor-derived ligands, with NKp46 binding to ecto-calreticulin on the surface of tumor cells [14], NKp30 recognizing B7-H6 [15] and HLA-B-associated transcript 3 (BAT3) [16], and NKp44 has been shown to bind to various tumor ligands, including proliferating cell nuclear antigen (PCNA) [17], platelet-derived growth factor D (PDGF-DD) [18], nidogen 1 [19] and HLA-DP [20]. With the notable exception of B7-H6, the relevance of the reported NCR ligands remains unclear.

Several therapeutic strategies focusing on NK cells have been conceptualized and are currently in various stages of development, from preclinical investigations to clinical trials [21]. However, the factors regulating NK cell-mediated tumor killing, including the variety of tumor ligands recognized by NK cells, but also the functional mechanisms employed by NK cells to kill tumor cells, are not yet fully understood. Here, we performed a genome-wide CRISPR/Cas9 screen to investigate the mechanisms of NK cell-mediated killing of the colon adenocarcinoma cell line HCT-116. Our findings reveal a critical involvement of tumor ICAM-1 in NKp44-dependent lysis of tumor cells. Furthermore, loss of factors within the apoptosis and IFN-γ signaling pathways conferred increased resistance of HCT-116 tumor cells to NK cell-mediated killing, highlighting the importance of these pathways in governing NK cell cytotoxicity. Additionally, our analyses demonstrate the direct antitumor role of NK cell-derived IFN-γ both *in vitro* and *in vivo*, further emphasizing the probable antitumor NK cell activity beyond the synapse formed between NK cells and their cognate targets.

## RESULTS

### A genome-wide CRISPR/Cas9 screen identifies tumor genes that modulate killing by NK cells *via* NKp44

We performed genome-wide CRISPR screens in HCT-116 colon carcinoma cells to identify functional perturbations affecting their killing by human KHYG-1 NK cell line through NKp44 engagement. The human NK cell line KHYG-1 does not express inhibitory KIR2DL1/L2/L3L4, nor KIR2DS1/S2 (**Figure S1**), and showed spontaneous cytolytic activity against HCT-116 cells (**Figure S2**). HCT-116 cells are sensitive to NK cell-mediated lysis, which is mainly induced by NKp44, as the frequency of killing is markedly reduced when an anti-NKp44 antibody is used (**Figure S2**). Invalidation of NKp44 in KHYG-1 cells also reduced the ability of NK cells to lyse HCT-116 cells. NKp44 can be encoded by 3 transcripts, with NKp44 T2 and NKp44 T3 encoding proteins with short cytoplasmic domains, while NKp44 T1 encodes a protein with an immunoreceptor tyrosine-based inhibitory motif (ITIM) that transmits an inhibitory signal to NK cells [22–24]. Transfection of all three transcripts encoding NKp44 was able to restore NKp44-mediated lysis (**Figure S3**). HCT-116 cells do not express PDGF-DD, NID1, HLA-DP or surface PCNA (**Figure S4**), which are ligands for NKp44 previously reported [25], suggesting that other NKp44 ligands are expressed at the cell surface and remain to be identified.

A clonal isolate of HCT-116 cells showing high activity of inducible Cas9 was generated (**Figure S5A**) and validated for its sensitivity to killing by NKp44 triggering (**Figure S5B**). Co-culture of HCT-116 Cas9^+^ cells were co-cultured with KHYG-1 effector cells allowed to determine that an effector-to-target ratio (E:T) of 2.5:1 exerts sufficient selection pressure for for the screening of target cell resistance to KHYG-1 killing in a 4 hour assay. Cas9-expressing HCT-116 cells were infected with Brunello’s genome-wide single guide RNA (sgRNA) library targeting 19,114 unique coding genes with 4 different sgRNAs per gene, and one thousand non-targeted controls. The mutant collection was co-cultured with KHYG-1 cells, KHYG-1 cells and anti-NKp44 antibody (α-NKp44 mAb) or NKp44-deficient KHYG-1 cells for 4 hours and cell death of KHYG-1 was promoted, a critical step that allowed the recovery of genomic DNA preparations that were not diluted by effector cell DNA (**Figure 1A** and **Material and Methods**). After 72 hours, a time frame that allows tumor target cell death by both direct NK cell cytolytic activities and the longer-term effects of cytokines released by the NK cells, tumor cells were recovered and subjected to a second round of KHYG-1 pressure, respectively (**Figure 1A**). The number of HCT-116 cells was reduced by 3.5, 2 and 1.8-fold at the end of the first round of selection and by 6.8, 3.1 and 1.9-fold at the end of screening in co-culture with KHYG-1 cells, KHYG-1 cells plus anti-NKp44 mAb or NKp44-deficient KHYG-1 cells, respectively (**Figure 1A**). Deep sequencing was further used to compare the changes in sgRNA abundance between the unchallenged state and the NKp44-independent states (KHYG-1 cells + α-NKp44 mAb, condition 2, **Table 2**; NKp44-deficient KHYG-1 cells, condition 3, **Table 3**) and the NKp44-dependent+independent state (KHYG-1 cells (condition 1, **Table 1**) (**Figure 1B**). One hundred and nineteen hits with a false discovery rate (FDR) of less than 0.5 were identified as significantly enriched in NKp44-dependent killing of HCT-116 cells by subtracting the hits obtained in conditions 2 and 3 from condition 1 (**Table 4**, **Figure 1** and **Material and Methods**). The enriched genes were then ranked using MAGeCK software (**Figure 1C** and **Table 4**).

**Figure 1:**
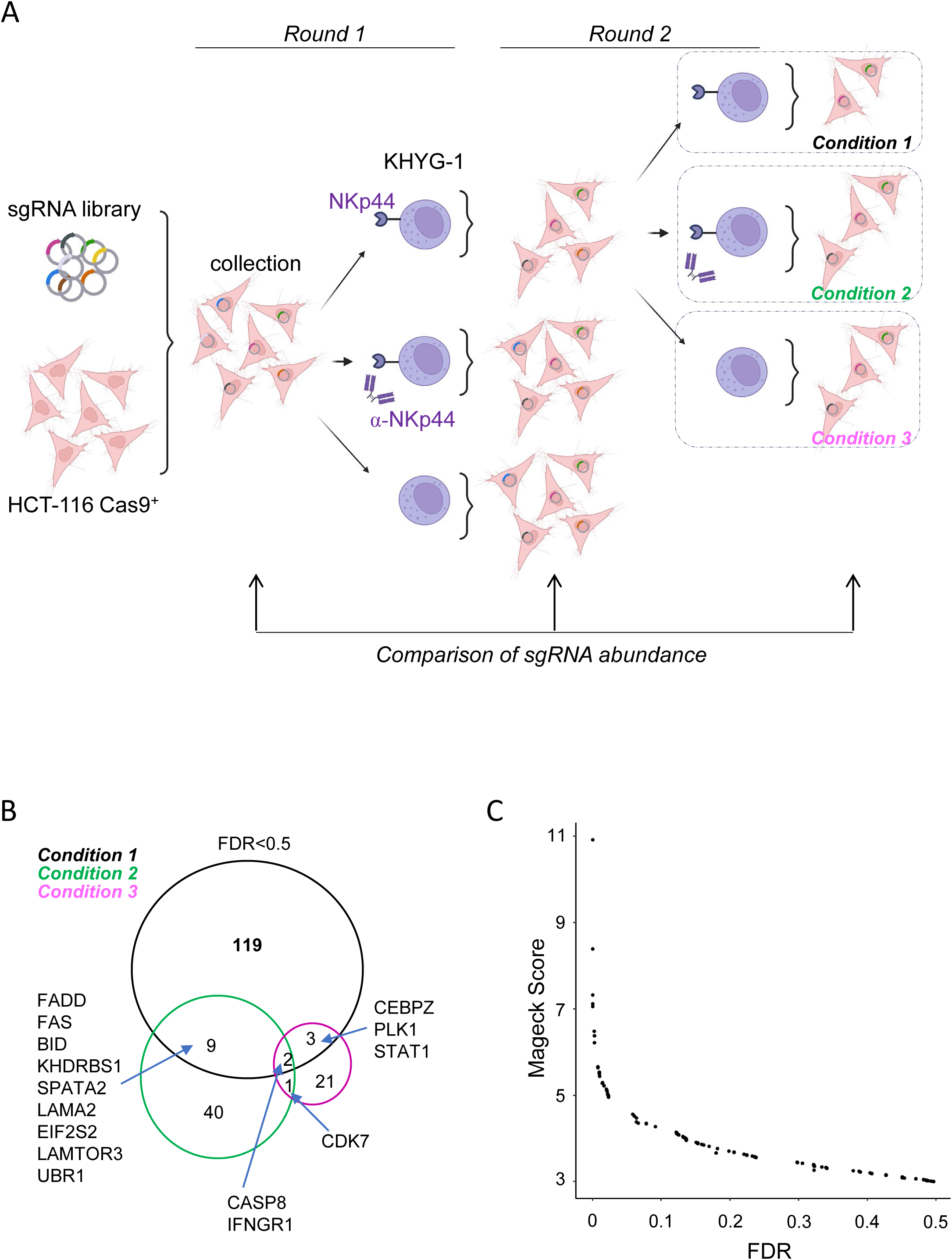
Genome-wide CRISPR/Cas9 screening identifies genes involved in NKp44-mediated specific killing of tumor cells by KHYG-1 cells. (A) Genome-wide CRISPR/Cas9 screening design. HCT-116 Cas9^+^ cells were transduced with the knock out sgRNA Brunello library. Collection of mutant cells were subjected to lysis by WT KHYG-1 or NKp44-deficient KHYG-1 in the presence of α-NKp44 mAb or not (Round 1). Mutant cells that survived lysis by KHYG-1 were subjected to a second round (Round 2) of co-culture with WT KHYG-1 (condition 1), in the presence of α-NKp44 mAb (condition 2) or with NKp44-deficient KHYG-1 (condition 3). The abundance of sgRNA in the collection, in the surviving cells in Round 1 and in the surviving cells in Round 2 were determined by sequencing (See Material & Methods). (B) Venn diagram of hits obtained with KHYG-1 co-culture (condition 1, See Table 1), KHYG-1+αNKp44mAb (condition 2, See Table 2) and NKp44-deficient KHYG-1 cells (condition 3, See Table 3). (B) Scatter plot showing the ranking of hits enriched in the NKp44-dependent killing of tumor cells by KHYG-1 cells by MAGecK score and false discovery rate (See Table 4).

### Critical role of ICAM-1 in NK cell lysis mediated by NKp44

To complete the dissection of the molecules involved in the synapse formed between NK cell and their cognate target cells, we first focused on the predicted cell surface proteins and identified 16 genes involved in the NKp44-dependent susceptibility of HCT-116 tumor cells to NK cell killing (**Figure 2A**). To assess these hits, we generated individual gene knockout (KO) cells by lentiviral sgRNA expression, which gave rise to polyclonal cell lines with varying degrees of gene disruption. This was determined by TIDE (Tracking of Indels by Decomposition) analysis, which allows to determine the spectrum and frequency of targeted mutations generated in a pool of cells by genome editing tools. We selected only mutant cell lines with an indel frequency (i.e. insertions and deletions) above 70%. Target KO and control cell lines were were subjected to KHYG-1-mediated lysis in the presence or absence of α-NKp44 mAb in a 4-hour chromium lysis assay (**Figure 2B and C**). Individual inactivation of the 16 target genes led to different levels of sensitivity to NK cell-mediated lysis (**Figure 2B**), but in each case addition of the α-NKp44 ab consistently inhibited the residual lysis suggesting that none of the inactivated target was a direct ligand of NKp44 (**Figure 2C**). We further conducted individual knockdowns of 103 out of the 119 hits from the screening with an FDR below 0.5, regardless of their predicted subcellular location, to assess their susceptibility to NKp44-mediated killing by NK cells. Additionally, we knocked down all genes encoding proteins thought to be expressed at the cell surface with an FDR of 0.74 (e.g., LTBR and PHLDA3). The lysis was inhibited to varying degrees by α-NKp44 mAb across all tested candidates (data not shown). We did not identify any gene knockout in HCT-116 cells that completely prevented the inhibition of lysis by the a α-NKp44 mAb. Notably, the least pronounced inhibition of lysis by α-NKp44 mAb was observed in tumor cells lacking the ICAM-1 gene (**Figure 2C**).

**Figure 2:**
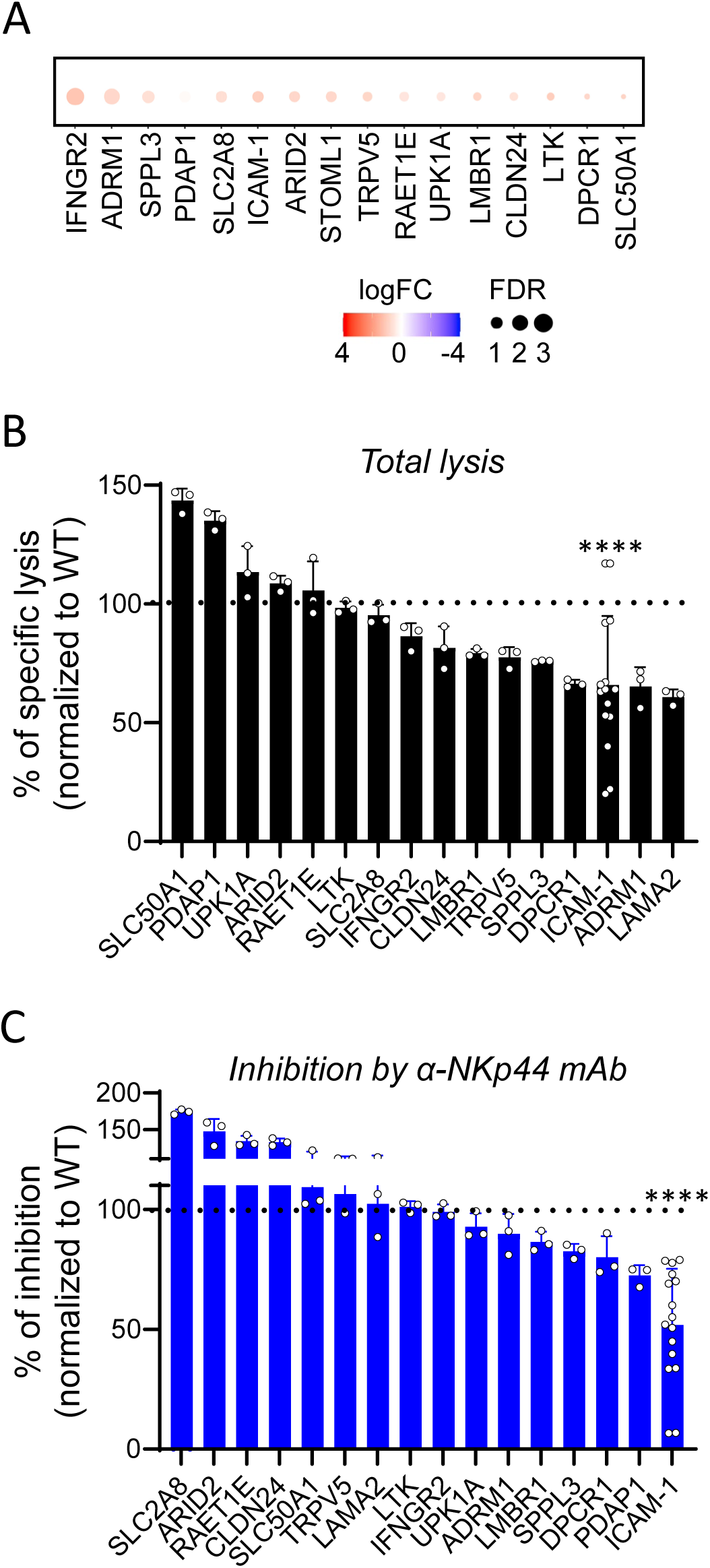
Knock-out of ICAM-1 in HCT-116 cells impairs NKp44-mediated lysis. (A) Enrichment of surface-encoding genes. (B-C) Indicated mutant cells were subjected to KHYG-1 killing in a standard 4-hour chromium release assay in the presence or not of α-NKp44 mAb. The frequencies of specific lysis normalized to WT were shown in (B) and the frequencies of inhibition by α-NKp44 mAb were shown in (C). *p* value<0.0001, unpaired t-test.

We then sought to dissect further the role of ICAM-1 expressed on tumor cells in their elimination by NK cells. LFA-1 is one of the ligands of ICAM-1 [26] and the crucial role of ICAM-1-LFA1 interactions in the initial adhesion of target lymphocytes and the polarization of cytotoxic granules towards the synapse is well described [27–30]. In the tumor context, although downregulation of ICAM-1 on acute myeloid leukemia cells in hematologic malignancies has been described to escape natural NK cell cytotoxicity [31], the role of ICAM-1 in solid tumors has not been fully investigated in terms of NK cell-mediated tumor killing. The ICAM-1 hit appeared to be significantly enriched upon NKp44-dependent lysis of HCT-116 cells (**Table 4**, **Figure 1C** and **2A**). Blocking or disrupting ICAM-1 reduced the total amount of killing by KHYG-1 cells (**Figure 2B**), but also significantly prevented NKp44-mediated killing (**Figure 2C**), arguing for either a key role of ICAM-1 in the NKp44-associated synapse or as a direct ligand for NKp44. We further sought to dissect further the role of ICAM-1 expressed on tumor cells in their elimination by NK cells (**Figure 3**). Increased ICAM-1 expression levels have been observed in multiple myeloma [32] and across many disparate carcinomas including breast [33], gastric [34], pancreas [35] and thyroid [36–38]. Sequencing the cDNA encoding ICAM-1 in HCT-116 cells revealed the expression of ICAM-1 transcripts #1 (including the LFA-1 binding domain) and #2 (lacking the LFA-1 binding domain), each with single nucleotide polymorphisms (SNPs) P352L and K469E in a heterozygous state (**Figure S6**). Expression of the wild-type (WT) or P352L/K469E SNP variants of the ICAM-1 transcript #1 in ICAM-deficient HCT-116 cells restored their susceptibility to NKp44-dependent killing by KHYG-1 cells (**Figure 3A**). ICAM-1 KO HCT-116 cells expressing the WT or P352L/K469E SNP variants of the ICAM-1 transcript #2, which lacks the binding domain for LFA-1 (**Figure S6**), did not restore the susceptibility to NK cell-mediated lysis as observed with transcript #1 (**Figure 3A**), arguing for a crucial role of NK cell LFA-1 in the ICAM-1-dependent elimination of HCT-116 cells by NK cells. We then investigated whether ICAM-1 transcripts could induce NKp44-dependent NK cell killing. HEK 293T cells, which do not express ICAM-1 (**Figure S6**), were resistant to NKp44-mediated killing by NK cells but remained sensitive to NKp30-mediated killing, confirming their overall susceptibility to NK cell-mediated cytotoxicity (**Figure 3B**). Expression of ICAM-1 transcript #1 in ICAM-1-deficient HEK 293T cells promoted target cell killing. However, this killing was not inhibited by α-NKp44 mAb (**Figure 3B**). To eliminate any confounding effects from ICAM-1/LFA-1 interactions, we knocked down LFA-1 in KHYG-1 cells (**Figure S7**). Expression of ICAM-1 did not promote the killing of HEK 293T cells by LFA-1 KO KHYG-1 cells, and the killing was not disrupted by the addition of α-NKp44 mAb (**Figure 3D**). In contrast, NKp30-mediated lysis of ICAM-1-negative HEK 293T cells (**Figure 3B**) and K562 cells in the presence of a blocking ICAM-1 antibody were observed (**Figure 3D** and **S6**). Taken together, these results suggested that ICAM-1 is required for NKp44-mediated HCT-116 tumor killing by NK cells, but is not sufficient on its own to promote HEK 293T killing *via* NKp44. However, ICAM-1 expression on other tumor cells from hematologic or solid origin may be dispensable for NK cell-mediated lysis by NKp30.

**Figure 3:**
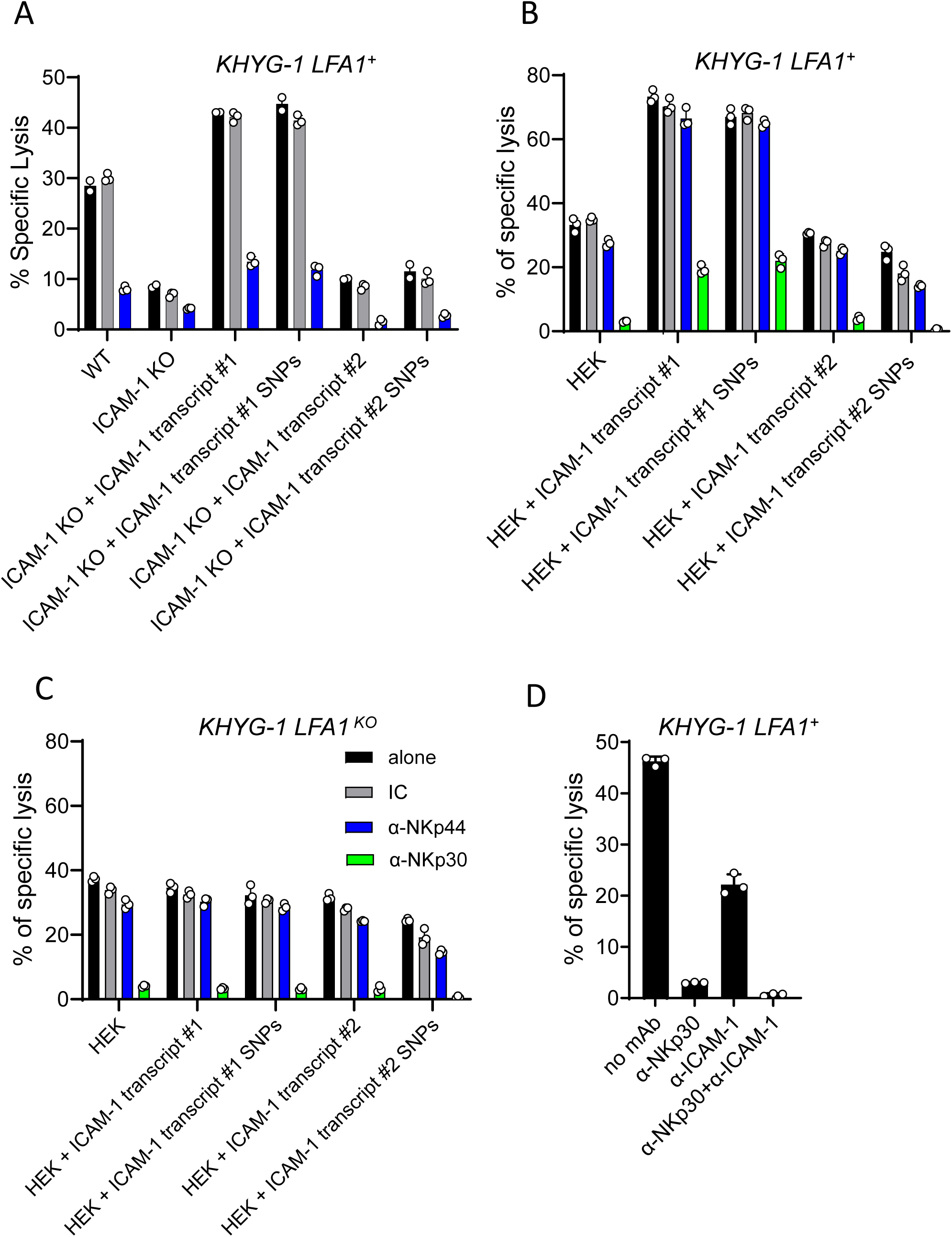
Expression of ICAM-1 is required but not sufficient for NKp44-mediated NK cell lysis of tumors. HCT-116 cells (A), ICAM-1-deficient HCT-116 cells (A), HEK 293T cells (B, C) and K562 cells (D) were cultured with KHYG-1 in the presence or not of α-NKp44, α-NKp30 or α-ICAM-1 mAbs as indicated. The frequencies of specific lysis were determined in a standard 4-hour chromium release assay. Data are representative of 2 to 3 independent experiments.

### Identification of tumor signaling pathways involved in sensitivity to NK cells

We then focused on the tumor genes influencing the sensitivity of HCT-116 tumor cells to cell-mediated lysis cells of KHYG-1 cells irrespective of NKp44 involvement (**Figure 4**). The top-ranked genes revealed by the screening include two broad classes of hits: sgRNAs targeting components of the apoptosis pathway and components of the IFN-γ signaling pathway, namely *FADD*, *CASP8*, *BID, FAS*, *FAF1* and *JAK1*, *IFNGR1*, *IFNGR2*, *JAK2* and *STAT1*, respectively (**Figure 4B-D**).

**Figure 4:**
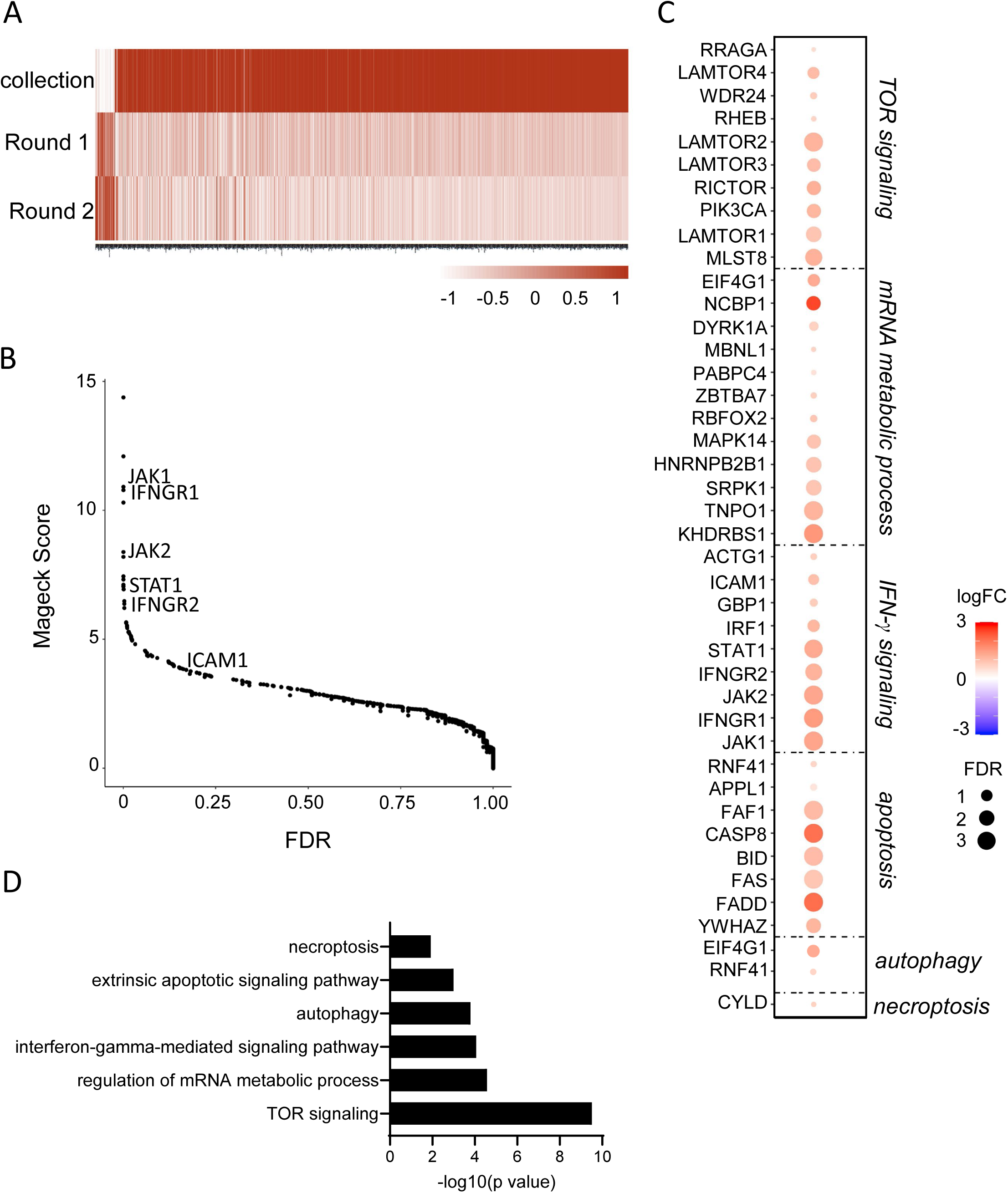
Identification of tumor signaling pathways involved in sensitivity to NK cells. (A) Heatmap of gene enrichment in rounds 1 and 2 in the KHYG-1+HCT-116 co-culture (condition 1 in Figure 1, See Table 1). (B) Scatter plot showing the ranking of hits enriched in the NK cell-dependent killing of tumor cells by KHYG-1 cells by MAGecK score and false discovery rate. (C-D). Selected enrichment of hits in the indicated pathways.

In addition, the screening revealed mechanisms of tumor cell sensitization to KHYG-1 cells that have not previously been associated with NK cell promotion of tumor cell death and require further investigation, with GO terms for mammalian target of rapamycin (mTOR) signaling, regulation of mRNA metabolic process, autophagy and necroptosis that were significantly enriched (**Figure 4C** and **D**).

The prominent role of apoptosis, both clinically and in our study, prompted us to compare the relative importance of perforin and FasL-mediated killing of HCT-116 tumor cells over time. We knocked down the expression of *PRF1* encoding perforin in KHYG-1 cells and that of *FAS* and *IFNGR1* (the ligand binding chain (alpha) of the gamma interferon receptor) in HCT-116 cells and assessed NK cell-mediated killing of target cells in a chromium release assay after 4 hours of co-culture (short time assay) and, as in our screening, 72 hours after co-culture (long time assay) by measuring the number of live tumor cells using an IncuCyte Imaging System (**Figure 5**). Killing of *FAS* KO target cells by KHYG-1 cells were ∼40% (SD ±14%) lower compared to wild-type HCT-116 cells, and in the absence of perforin, *FAS* KO cells were not killed by KHYG-1 effectors (**Figure 5A**). These results suggested that the direct cytolytic effect on HCT-116 cells after a 4-hour co-culture with KHYG-1 cells mainly required perforin expression by NK cells and depended to a lesser extent on Fas-mediated apoptosis. Of note, killing of K562 leukemia cells, a prototypical NK cell target, was completely dependent on perforin expression by KHYG-1 cells after 4 hours of co-culture (not shown). Seventy two hours after co-culture with NK cells, the number of remaining *FAS* KO cells was ∼65% (SD ±17%) lower than that of WT target cells, suggesting that the Fas/FasL interaction was the main mechanism for tumor cell death at this later time point (**Figure 5B**). The perforin-dependent mechanism promoting tumor cell death was less pronounced at a later time point, as the frequency of remaining cells was ∼20% (SD ±10%) lower when cells were co-cultured with perforin KO KHYG-1 cells (**Figure 5B**). Taken together, these results suggested that apoptosis is a key mechanism in NK cell-mediated killing of HCT-116 tumor cells. Initially, after encountering the tumor cells, this process is largely dependent on perforin, but later on the interaction between Fas and FasL.

**Figure 5:**
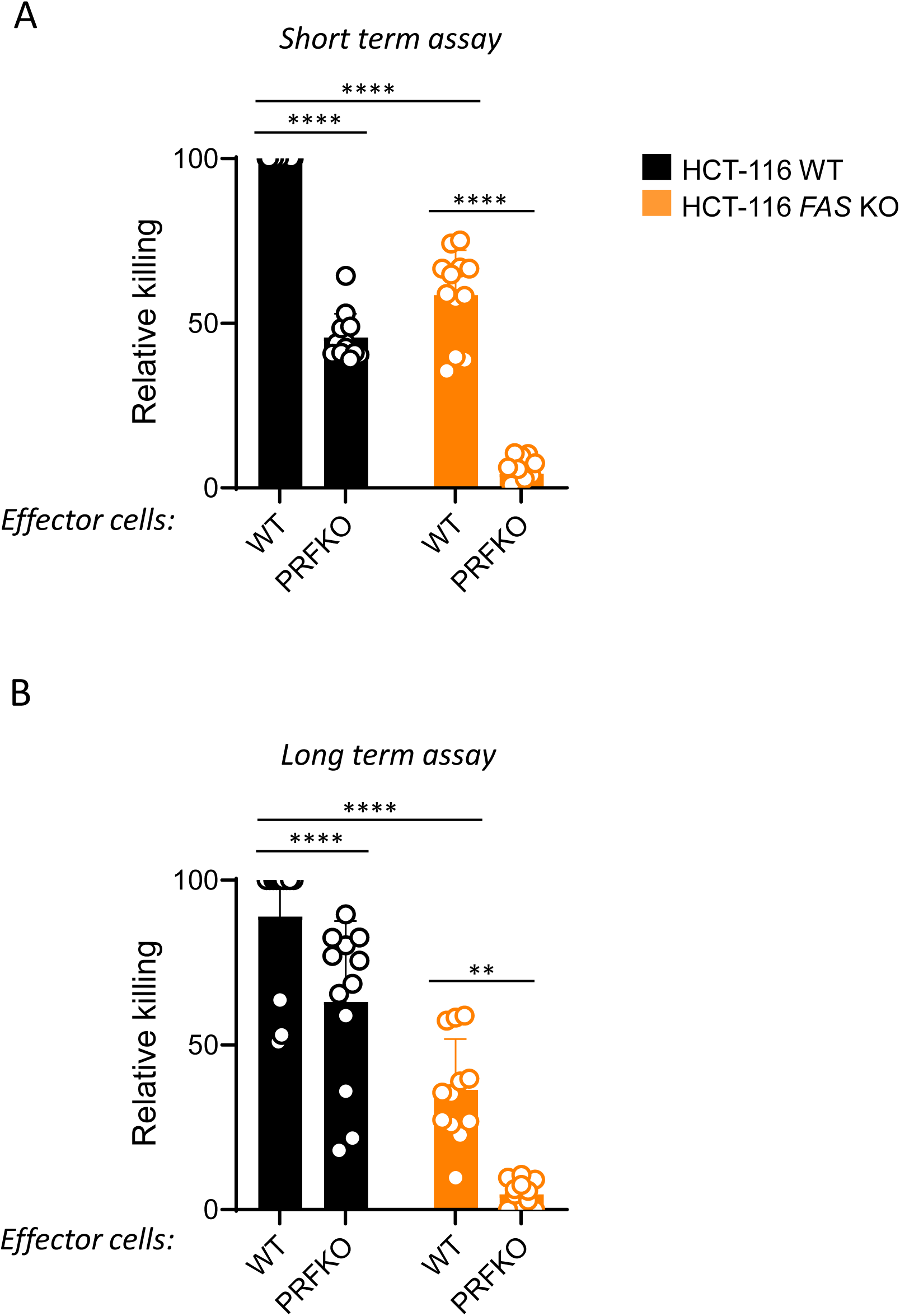
Relative requirement of NK cell effector functions for NK cell -mediated tumor lysis over time. WT and Fas-deficient HCT-116 Cas9^+^ cells were co-cultured with WT KHYG-1 or perforin-deficient KHYG-1 cells. The extent of specific lysis was determined in a standard 4-hour chromium release assay (short term) (A) or as the CRISPR screen condition (long term) by estimating the number of live cells with an Incucyte Imaging System (B). P values: ****<0.0001; **=0.0037. Unpaired T test. Data result from the pool of 3 independent experiments.

### IFN-γ produced by NK cells/ILC1s is critical for the control of B16F10 melanoma in a mouse model of disseminated disease

Our results showed that mutations in the interferon-γ receptor (IFNγR) signaling pathway make HCT-116 tumors more resistant to destruction by activated NK cells. We thus assessed *in vivo* the requirement for NK cell-derived IFN-γ in the control of metastasis in a mouse model of B16F10 melanoma. The control of B16 melanoma cells by NK cells and/or type 1 innate lymphoid cells (ILC1) *in vivo* is well-documented [39], with NK cells and ILC1 sharing phenotypic characteristics, including the expression of NK1.1 and NKp46 surface markers [40, 41]. We observed a marked increase in metastatic foci in the lungs of mice treated with a depleting anti-NK1.1 mAb and in mice deficient in NKp46^+^ cells (*Ncr1^iCre^*^/+^*R26^DTA^* mice), with ∼11 (SD ±5) and 4 (SD ±1) times more metastatic foci, respectively, compared to controls (**Figure 6**, upper panel). Additionally, treatment of wild-type (WT) mice with a blocking anti-IFN-γ mAb resulted in a ∼2.5-fold (SD ±1) increase in lung metastases (**Figure 6**, bottom panel). Remarkably, mice with NKp46^+^ cells knocked down for *Ifng* (*Ncr1^iCre^*^/+^*Ifng^flx/flx^* mice) exhibited a substantial increase in metastases compared to control mice (*Ncr1^iCre^*^/+^*Ifng^+/+^* mice). In these knockout mice, *Ifng* deletion occurs specifically in NKp46^+^ cells, including NK cells, ILC1s, and NKp46^+^ ILC3s, but not in CD8^+^ T cells or NKT cells (**Figure S8**, [42] and not shown). These results collectively demonstrated that IFN-γ produced by NKp46^+^ cells is crucial for the control of tumor cell metastasis *in vivo*.

**Figure 6:**
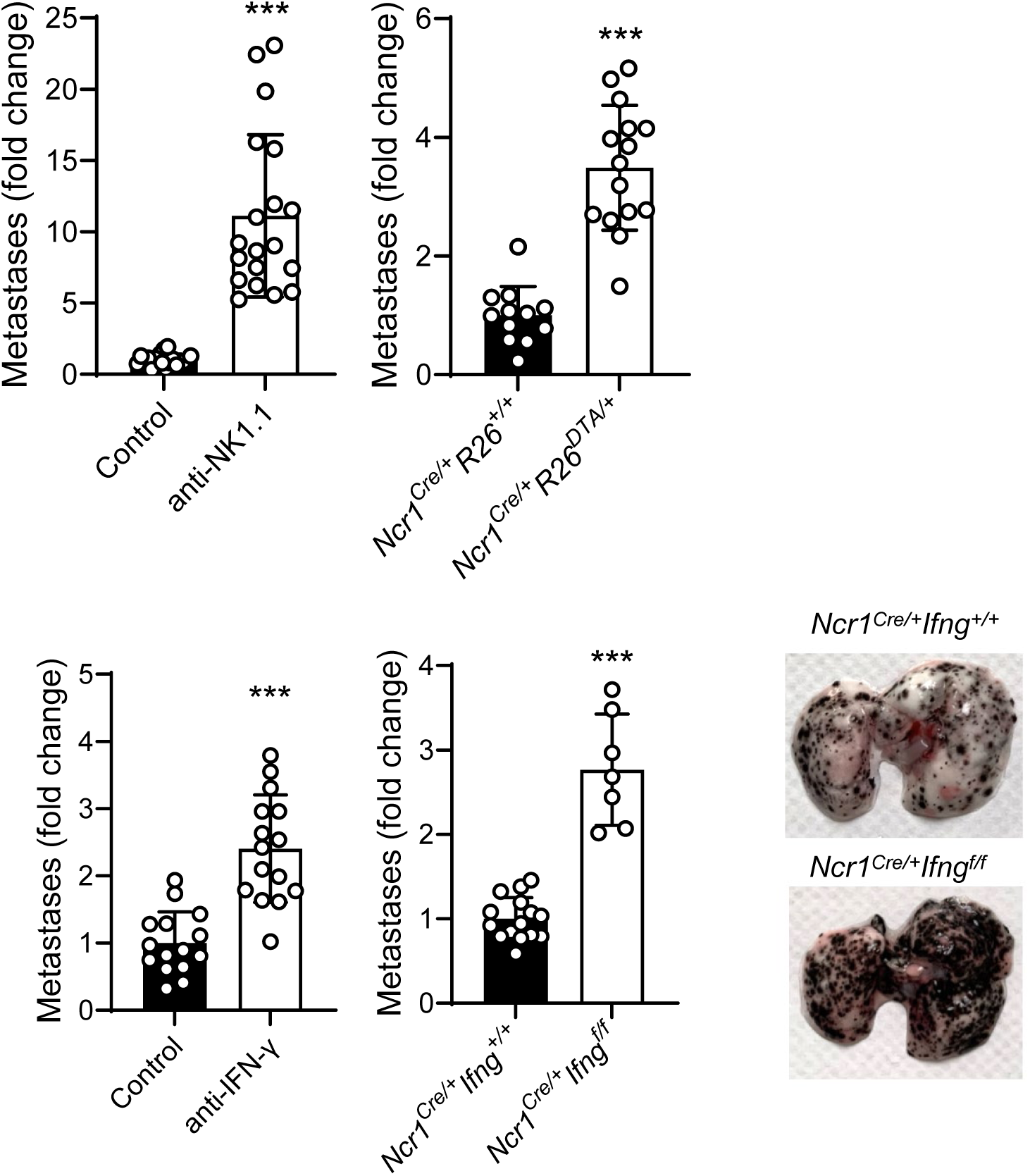
IFN-γ from NKp46^+^ cells is required to control the metastasis of B16F10 in the lung. WT mice treated with depleting α-NK1.1 mAb or with neutralizing IFN-γ mAb, *Ncr1^iCre/+^R26^DTA/+^*mice, *Ncr1^iCre/+^Ifng^fl/fl^*) and their respective controls (mice treated with isotype controls, *Ncr1^iCre/+^R26^+/+^* or *Ncr1^iCre/+^Ifng^+/+^*) were injected intravenously with B16F10 cells. The number of foci in the lungs were determined 7 days later. Data represent 1 to 4 independent experiments; mean ± SD; Student test with Welsh correction ***p<0.005.

### IFN-γ produced by NK cells promotes mitochondrial apoptosis in tumor cells

The requirement of IFN-γ from NK cells for tumor control in vivo may be indirect, by mobilizing other immune system players, or may act directly on the tumor cells themselves. IFN-γ can stimulate the activation of JAK/STAT, which in turn drives apoptosis directly or through induced lipid peroxidation and ferroptosis [43]. [43]. We thus assessed if IFN-γ can directly promote tumor cell death. The number of live WT HCT-116 cells was reduced by 26% (SD ±10%) and 46% (SD ±4%) when treated with 1 and 10 μg/mL recombinant human IFN-γ, respectively, compared to IFNGR1 KO cells, highlighting the antitumor role of IFN-γ (**Figure 7A**). We investigated whether IFN-γ was also able to promote tumor cell death by inducing the mitochondrial-mediated apoptosis pathway, a described NK cell-mediated tumor killing mechanism. WT and IFNGR1 KO HCT-116 cells were co-cultured with KHYG-1 cells and the amount of cytochrome C released in the cytosol was monitored by FACS (**Figure 7B**). The frequency of IFNGR1 KO cells that had released their cytochrome C in the cytosol was ∼44% lower compared to WT HCT-116 cells (**Figure 7B**), suggesting that NK cell-derived IFN-γ promotes tumor killing in part through induction of the mitochondrial-mediated apoptosis pathway.

**Figure 7:**
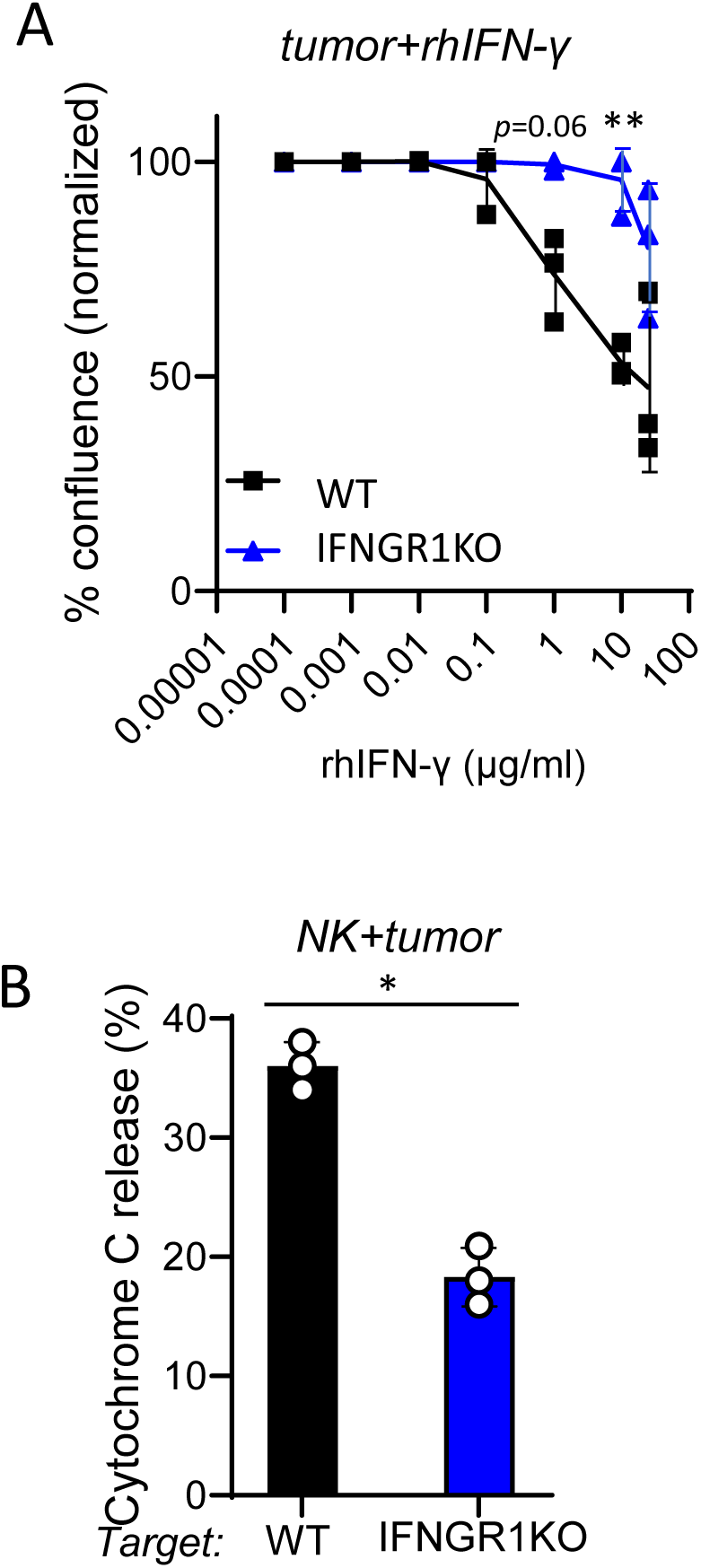
IFN-γ can directly kill tumor cells in part by triggering mitochondrial apoptosis. (A) WT or IFNGR1 KO HCT-116 cells were cultured in the presence of the indicated amount of recombinant IFN-γ for 72 hours. The number of live tumor cells was estimated using an Incucyte Imaging System. Two-way anova with Dunnett’s comparison test. (B) KHYG-1 cells were co-cultured with WT or IFNGR1 KO HCT-116 cells for 24 hours. The amount of cytochrome C released in target cells were determined by FACS. Data are representative of 2 independent experiments. mean ± SD; Student test.

## DISCUSSION

Understanding how tumor-intrinsic factors influence the mechanisms of NK cell-mediated killing is crucial for advancing NK cell-based anti-tumor therapies. In our study, we performed CRISPR/Cas9 screens using KHYG-1 NK cells and HCT-116 tumor cell co-cultures to identify genetic perturbations that impact NK cell-mediated immunity against cancer. One screen was specifically designed to explore NK cell-mediated tumor cell killing via NKp44. We identified 119 genes with a false discovery rate of less than 0.5, which were significantly enriched in the NKp44-dependent killing of HCT-116 cells by KHYG-1 cells. These genes should include those involved in the signaling pathways necessary for target cell killing by KHYG-1, as well as pathways that regulate the expression of NKp44 ligands, including the ligand itself. However, we were unable to pinpoint genes whose invalidation specifically inhibits tumor cell killing through NKp44. Several hypotheses might explain this result. One possibility is that HCT-116 cells co-express multiple ligands for NKp44. If this is the case, knocking out a single ligand would still permit KHYG-1 cells to kill tumor cells via NKp44 through interaction with the remaining ligand(s). Alternatively, the NKp44 ligand might be essential for HCT-116 cell survival. Although CRISPR/Cas9 technology allows genome-wide screening, it limits analysis of cells mutated for essential genes. In our screening with the CRISPR knockout sgRNA library Brunello, approximately 400 mutant cells with survival gene knockouts were lost after genome editing, corresponding to an expected loss of 2.6% of genetic information in such experiments [44]. If the genes regulating NKp44 ligand expression are linked to survival genes, we could not identify them using this technology.

Despite these limitations, our study found that disruption of ICAM-1 significantly reduced NKp44-mediated lysis of HCT-116 cells by KHYG-1 cells, more than disruption of other candidate genes. ICAM-1 expression is typically low under non-inflammatory conditions and mainly detectable on endothelial cells. However, upregulation of ICAM-1 is a common response to inflammatory stimuli, including pro-inflammatory cytokines, viral infections [45], and cancer [32–38]. ICAM-1 serves as a cellular entry point for major group rhinoviruses [46], certain coxsackieviruses [47], and various bacterial pathogens [48]. The increased expression of ICAM-1 on the surface of infected cells enhances infectivity [49] but also likely signals the immune system to target these cells for destruction by promoting leukocyte adhesion, thus facilitating an immune response. ICAM-1 may itself act as a stress ligand for immune cells. Notably, ICAM-1 is the tumor antigen targeted by CAR T cells in novel treatments currently being evaluated for thyroid cancer patients [50]. Given the ability of NK cells to sense cellular stress [21], we investigated the role of ICAM-1 as a potential stress ligand for NKp44. Our results demonstrated that ICAM-1 expression is required for NK cell-mediated killing of HCT-116 targets via NKp44, as restoring ICAM-1 expression rescued the killing of ICAM-1-deficient HCT-116 cells. However, expressing ICAM-1 in a tumor cell line that is inherently ICAM-1 deficient and resistant to NKp44-mediated lysis was insufficient to promote lysis. These findings suggest that ICAM-1 is not a direct ligand for NKp44. Several explanations might account for this result. One possibility is that ICAM-1 multimerization is required for effective binding to NKp44. Supporting this hypothesis, we identified the SNP variant K469E in ICAM-1 transcripts from HCT-116 cells, which may promote the formation of higher-order ICAM-1 assemblies on the cell surface [51]. Artificial expression of ICAM-1 in HEK 293T cells might not accurately replicate the natural cell surface expression necessary for NKp44 binding in HCT-116 cells. Furthermore, ICAM-1 is a highly glycosylated protein with eight N-glycosylation sites, and the extent of glycosylation is known to regulate ligand binding [52] playing a crucial role in modulating ICAM-1’s role [53]. Differences in the glycosylation profiles of ICAM-1 in HCT-116 tumors compared to those in HEK 293T cells might prevent NKp44 binding, a possibility that remains to be investigated. Finally, the co-engagement of other receptors on NK cells might be necessary to induce lysis. While the interaction between ICAM-1 and NKp44 may be sufficient for the lysis of HCT-116 cells, it might not be enough to influence HEK 293T cell lysis. ICAM LFA interaction may fascilitate Nkp44 clustering at the synapse and/or stabilize NKp44 interaction with a low affinity ligand. Nevertheless, it remains the case that ICAM-1 is a crucial component of NKp44-dependent NK cell killing. Thus, the necessity of ICAM-1 may vary depending on the other molecular partners that form the synapse, reflecting the complex diversity of NK cell-dependent tumor cell recognition.

The interaction between LFA-1 and ICAM-1 is crucial for the formation of the immunological synapse and the cytolytic activity of T cells, where LFA-1 co-stimulation plays a crucial role in granule polarization and degranulation [29, 54]. In resting NK cells, LFA-1 signaling induces granule polarization, while degranulation is triggered by activating receptors independently of LFA-1 [55–58]. Our results indicate that ICAM-1 expression is necessary for NK cell-mediated killing of HCT-116 tumor cells. However, the LFA-1/ICAM-1 interaction is not essential for the killing of K562 and HEK 293T cells, suggesting that the requirement for this interaction can vary depending on the specific tumor target. Other integrin pairs on tumor and effector NK cells may compensate for the absence of LFA-1/ICAM-1 interaction in these cases. Moreover, our assays utilized IL-2-activated KHYG-1 cells, which may depend less on LFA-1/ICAM-1 interactions for degranulation compared to resting NK cells. This suggests that the activation status of NK cells can influence their reliance on LFA-1/ICAM-1 interactions. However, it has been shown that ICAM-1 expression on tumor cells is crucial for antibody-dependent cellular cytotoxicity mediated by CD16 on IL-2-activated primary NK cells [59]. Therefore, monitoring or promoting ICAM-1 expression on tumor cell surfaces should be considered in the context of NK cell-based therapies.

Our study also reveals that mutations in the interferon-γ receptor (IFNγR) signaling pathway, specifically in genes such as *IFNGR1*, *IFNGR2, STAT1, JAK1*, or *JAK2*, render HCT-116 tumors more resistant to killing by activated NK cells. The critical role of the IFN-γR pathway in determining tumor cell susceptibility [60] or resistance [61–63] to NK cells underscores the complexity of IFN-γ signaling in cancer. This complexity involves various mechanisms, including major histocompatibility complex (MHC) upregulation [30, 62, 63], growth suppression, and immune modulation. Recent studies have shown that IFN-γ can enhance NK cell-mediated tumor killing by upregulating adhesion molecules such as ICAM-1 on tumor cells [64]. Nevertheless, IFN-γ is a pleiotropic cytokine with direct antitumor functions. IFN-γ can activate the IDO1-Kyn-AhR-p27 pathway to induce tumor quiescence [65] and stimulate the p16INK4a-Rb pathway, triggering tumor senescence [66]. Additionally, IFN-γ stimulates the JAK/STAT signaling pathway, leading to increased caspase activity and a decrease in SLC7A11 and SLC3A2 [43], promoting apoptosis directly or through lipid peroxidation and ferroptosis [67]. Recently, IFN-γ derived from CD4^+^ CAR T cells has been identified as a key mechanism promoting apoptosis in tumor cells [68]. While NK cells have long been known to induce apoptosis, recent research highlights the importance of mitochondria-mediated apoptosis in NK cell-mediated tumor cell killing [69]. Granzyme B is a well-known promoter of this pathway, but our findings suggest that IFN-γ can also serve as an NK cell-derived mediator that activates mitochondria-mediated apoptosis, leading to tumor cell death. Notably, IFN-γ produced by CD4^+^ CAR T cells can induce apoptosis without direct cell contact. Given the significance of IFN-γ diffusion in bystander tumor cell killing by T cells [70, 71], it is plausible that NK cells substantially contribute to creating extensive cytokine fields within the tumor microenvironment. This process could enable the long-distance elimination of IFN-γ-sensitive tumor cells. Further investigation is needed to determine the extent to which this mechanism contributes to NK cell-mediated tumor cell killing.

*In vivo* studies support the role of NK cells and IFN-γ in tumor sensitivity. However, the specific contribution of NK cell-derived IFN-γ to tumor cell killing remained to be evaluated. RAG^-/-^ mice, which lack T and B cells, are resistant to B16 tumor cell growth arguing for a role of innate immunity [72]. This resistance is not observed in mice treated with anti-NK1.1 monoclonal antibody (mAb) or polyclonal anti-asialo-GM1 antibody, which deplete NK cells but also affect NKT cells (anti-NK1.1 mAb) [73] and γδ T cells (anti-aGM1 mAb) in the liver. Mice deficient in IFN-γ, the IFN-γ receptor, or components of the IFN-γ signaling pathway (e.g., *Stat1*^-/-^ mice) develop more MCA-induced tumors compared to wild-type mice [74, 75]. Using a mouse model lacking NKp46^+^ cells (*Ncr1^iCre^*^/+^*R26^DTA^* mice)-which include NK cells, ILC1s, and some γδ T cells-we demonstrated that NKp46^+^ cells are essential for controlling B16 metastasis formation in the lungs. Importantly, utilizing a unique mouse model of IFN-γ deficiency in NKp46^+^ cells, we showed that IFN-γ derived from NK cells and ILC1s is critical for controlling B16 metastasis in the lungs. Supporting the role of NK cell-derived IFN-γ in metastasis control, IFN-γ production by NK cells following infection with engineered *Salmonella typhimurium* was identified as the mechanism effectively suppressing metastasis across a range of cancers [76]. The direct impact of NK cell-derived IFN-γ on tumor cell growth control and its indirect effects through immune response promotion remain to be fully explored. Nevertheless, a higher number of IFN-γ-producing NK cells is associated with a favorable response to treatment for chronic myeloid leukemia [77]. In addition, more than one-third of melanoma and lung adenocarcinoma cell lines possess mutations in components of the IFN-γ signaling pathway [74]. Furthermore, mice bearing melanoma tumors with knocked-out IFNGR1 exhibit impaired tumor rejection after anti-CTLA-4 therapy [78]. Melanoma patients who do not respond to anti-CTLA-4 (ipilimumab) [78] or those with metastatic melanoma experiencing relapses after anti-PD-1 therapy (pembrolizumab) [79] have tumors with genomic defects in IFN-γ pathway genes. These defects result in a lack of response to IFN-γ, including insensitivity to its antiproliferative effects on cancer cells. However, studies indicate the deleterious effects of IFN-γ, which can induce the expression of immune checkpoint molecules such as PD-L1 on tumor cells [80], potentially inhibiting T cells. This dual effect of IFN-γ highlights its complex role in cancer therapy, balancing anti-tumor immunity and potential immune evasion mechanisms.

In conclusion, our findings encourage further research into how NK cell-derived IFN-γ can be harnessed to enhance direct tumor cell killing and investigate its bystander effects through cytokine diffusion. It is crucial to explore ways to leverage the benefits of IFN-γ while mitigating its potential drawbacks. Additionally, monitoring ICAM-1 levels on tumor cell surfaces or modulating its expression in NK cell-based therapies warrants consideration.

## Supporting information

reagents

## ACKNOWLEDGMENTS

We thank the CRISPR Screen Labtech, label Aix-Marseille Platform, of the CIML for reagents and advices. We thank the members of the CB2M (Computational Biology, Biostatistics and Modeling) group at CIML for their expert help with data analysis. We thank Daphné Jaubert for her participation in the *in vivo* experiments.

## CONTRIBUTORS

S.G., A.F., J.G., H.M-K, N.L., M.V., A.P., and E.N-M. performed the experiments. S.G. generated and selected the Cas9-expressing HCT-116 clone, amplified the Brunello library, generated the lentiviral particules and titrated the production on HCT-116 cells. S.G. and A.F. set up screening conditions. S.G., A.F., and E.N-M. conducted the genome-wide CRISPR/Cas9 screenings. S.G. prepared the libraries for sequencing. S.G., A.F., J.G., and H.M-K generated and tested the individual gene knockouts in HCT-116 using chromium release assays. A.F., with assistance from S.G., validated gene knockouts by flow cytometry and/or TIDE analysis. A.F. developed the KHYG-1 NKp44 knockout, LFA-1 knockout, and ICAM-1 transfectants, and conducted their analysis. S.G. and A.F. investigated roles of perforin, Fas, and IFN-γ, with assistance from A.P. S.G., A.F., and N.L. studied the role of IFN-γ in promoting cell death and cytochrome C release. S.G. and M.V. conducted the *in vivo* experiments. S.G. took care of the mice breeding and genotyping. A.P. has performed the KIR phenotyping. A.M. and S.R. provided the lentiviral plasmid, the Brunello library, advice for the screenings, the high throughput CRISPR/Cas9 KO protocol and bioinformatic analysis. B.E. performed the bioinformatic analysis. L.G. and B.R. provided advices on the biochemistry of NKp44. E.N-M. and E.V. conceived the project. E.N-M. wrote the manuscript with contributions from E.V. S.G and A.F. reviewed the manuscript. S.G and A.F. wrote the Materials and Methods section of the manuscript.

## PATIENT CONSENT FOR PUBLICATION

Not applicable.

## ETHICS APPROVAL

All mouse experiments were performed in accordance with the rules of the CIML ethics committee and were approved by the *Ministère de l’Enseignement Supérieur, de la Recherche et de l’Innovation* – France (APAFIS#30510-2021031817315100 v5).

## AVAILABILITY OF DATA AND MATERIAL

Data are available on reasonable request. All data relevant to the study are included in the article or uploaded as online supplemental information. The data that support this study are available from the corresponding author on reasonable request. Hits from the CRISPR/Cas9 KO screenings discussed in this publication have been deposited in NCBI’s Gene Expression Omnibus and are accessible through accession number #XXX.

## MATERIALS AND METHODS

### Cell lines

All cell lines were purchased from ATCC and tested weekly for mycoplasma contamination using the MycoAlert Mycoplasma Detection Kit (Lonza). All cell lines were cultured in RPMI 1640 medium (Gibco) supplemented with 10% heat-inactivated fetal bovine serum (FBS, Gibco), 2 mM L-glutamine (Gibco), 1% non-essential amino acids (Gibco), 1 mM sodium pyruvate (Gibco), 1% Hepes 1M (Gibco), and 1% penicillin/streptomycin (Gibco), and maintained at 37°C in an atmosphere containing 5% CO_2_. KHYG-1 cells were supplemented with 100 U/ml human IL-2 (Miltenyi ref#130-097-743). HEK 293T and HEK 293FT were cultured in DMEM medium (Gibco) supplemented with 10% heat-inactivated fetal bovine serum (FBS, Gibco), 2 mM L-glutamine (Gibco), 1% non-essential amino acids (Gibco), 1 mM sodium pyruvate (Gibco), 1% Hepes 1M (Gibco) and 1% penicillin/streptomycin (Gibco) and maintained at 37 °C in an atmosphere containing 5% CO_2_.

### Generation and selection of Cas9-expressing HCT-116 clones

For each transduction, 500,000 HCT-116 cells were plated in 12-well plates in 1 mL complete medium containing lentiviruses supplemented with 8 µg/mL polybrene (Sigma H9268) and cells were spun down at 950 x g for 90 min at 30°C. The cells were then maintained at 37°C in an atmosphere containing 5% CO_2_.

pCW-Cas9 blast and pLKO-based sgRNA vector were purchased from Addgene (plasmid #83481 and#52628, respectively). The puromycin gene was removed and replaced with a puro-GFP fusion protein previously described by Gibson assembly to obtain the pLKO1.5-puro-GFP vector (Phelan, Young et al. 2018).

A fraction of single cell clones was transduced with pLKO1.5-puro-GFP targeting RPL6, an essential gene for cell survival in human cells. Cas9 expression was induced by the addition of 200 ng/mL doxycycline (Sigma D3072) and the most efficient clones were selected based on the highest and fastest decrease in GFP+ cells among total cells by flow cytometry [81, 82].

### CRISPR Library

Human Brunello CRISPR knockout pooled library was provided by the CRISPR Screen LabTech platform (Addgene #73178).

### Virus production and transduction

See [81].

### Collection of genome-wide HCT-116 mutant cells

For genome-wide screens, Cas9-expressing HCT-116 cells were transduced such that an average of 500 copies of each sgRNA were present after selection. Cultures were continued for the duration of the 9-day screen while maintaining 500-fold coverage. The collection of mutant human HCT-116 cells was obtained by transducing 120 million HCT-116-Cas9+ cells with lentiviruses containing the Brunello library at an infection multiplicity of 0.3. The transduced cells were further cultured for 3 days under antibiotic selection pressure (2 μg/mL puromycin (Thermo Fisher)) until the non-transduced control cells were dead. Induction of Cas9 expression was initiated by addition of doxycycline (200 ng/mL) on day 4 after transduction. The transduced cells were counted and passaged every two days with fresh doxycycline-containing medium until the cells were harvested again on day 9 for DNA extraction.

### Genome-wide CRISPR/Cas9 screenings coupled to NK cell cytotoxicity

50 million genome-wide HCT-116-Cas9+ mutant cells were co-cultured with 125 million WT KHYG-1 NK cells with or without anti-NKp44 (10 µg/mL) or with NKp44-deficient KHYG-1 cells at an effector:target (E:T) ratio of 2.5:1. For the co-culture, 30,000 target cells and 75,000 effector cells were distributed in a 96-well flat bottom plate. After 4 hours, the cells were harvested and washed with PBS 2mM EDTA. Cells were then incubated for 72 hours in complete medium containing 2 µg/mL puromycin to eliminate KHYG-1 cells. 50 million of the surviving HCT116 mutant cells were frozen for subsequent DNA extraction and the remaining cells that survived in the condition with KHYG-1 (condition 1, Figure 1) were cultured until 150 million cells were reached to allow the second round of co-culture under the 3 different conditions. The second round was performed as described above and cells were frozen for DNA extraction. All cell pellets were washed twice with ice-cold PBS 1X and dry pellets were stored at −80°C.

### CRISPR library preparation for next-generation sequencing

For each sample, genomic DNA was isolated from the cryopreserved cell pellet using the DNeasy Blood and Tissue Kit (Qiagen) and the genomic DNA concentration was determined using a NanoDrop spectrophotometer. Library preparation requires a two-step PCR amplification, purification and quantification according to Webster et al. 2019. In a first PCR step, the sgRNA sequences are amplified with the PCR primers U6-F1 and Tracr-R1. A second PCR was performed to attach Illumina-compatible sequencing adapters for the NextSeq2000 sequencer (genomic platform at CIML). After purification on a 2% size-selective E-gel, the secondary PCR product was quantified using the Qubit dsDNA HS Assay Kit (Thermo Fisher) and qualified on the bioanalyzer (Agilent) using the High sensitivity DNA Assay (Agilent).

### Next-generation sequencing (NGS) and analysis

The pooled libraries were sequenced on an Illumina NextSeq2000 sequencer using a single-read P2 flow cell (550 million reads). Libraries were sequenced at a depth of ∼20 million reads per condition. Reads were aligned to the sgRNA library using Bowtie2. From these alignments, Protospacer count tables were generated using Python scripts and processed with MAGeCK. MAGeCK analysis was used to score and prioritize sgRNAs using the default settings of the algorithm (Li et al., 2014). MAGeCK scores were normalized to -log10 and values were plotted against FDR values.

### Generation of individual gene KO in HCT-116-Cas9^+^ cells

The sgRNA targeting each gene to be invalidate was acquired from IDT in a 5’ forward and 3’ reverse oligo format. The 5’ forward sgRNA and 3’ reverse sgRNA contain homologous sequences to BFUA1 enzyme to enable cloning into pLKO1.5-puro-GFP digested with BFUA1 enzyme. After annealing the forward and reverse oligos, the double-stranded DNA sgRNA were individually cloned into the pLKO1.5-puro-GFP vector using T4 DNA ligase (New England Biolabs), and the plasmids were transformed into chemically competent Stbl3 *E.coli* (Thermo Fisher). The bacteria were plated on petri dishes containing LB agar and 100 µg/mL ampicillin and cultured overnight at 37°C. The colony was harvested and the bacteria were cultured in LB medium containing 100 µg/mL ampicillin for 24 hours at 37°C and miniprep processed. The plasmid DNAs were sequenced to check whether each sgRNA was integrated into the plasmid in a good reading frame. Subsequently, lentiviral particles were generated using HEK 293FT cells as described in [81]. The transduction of HCT-116-Cas9+ with the lentiviral particules have been performed in p96W flat plate. For each condition, 100000 HCT-116-Cas9+ cells were plated in 96-Well plate in a final volume of 150µL of complete RPMI supplemented with 8µg/mL of polybrene (Sigma H9268). Cells were spun down at 950g for 90 min at 30°C. The cells were then maintained at 37°C in an atmosphere containing 5% CO_2_ for 3 days. The transduced cells were further cultured for 3 days under antibiotic selection pressure (4 μg/mL puromycin (Thermo Fisher)). Induction of Cas9 expression was initiated by addition of doxycycline (200 ng/mL) on day 6 after transduction. DNA extraction and cytotoxicity assay were performed at day 9 post transduction. Cells were frozen in 96-Well V shape with 30µL of Cryostor CS10 (StemCell #100-1061).

### Generation of NKp44, perforin and LFA-1 KO in KHYG-1 cells

Invalidation of NKp44, perforin and LFA-1 were achieved by nucleo-infection using a complex of Cas9 protein (IDT) and sgRNA targeting each individual target gene on the Neon machine using the Invitrogen™ Neon™ Transfection System 100 μL Kit. For the invalidation of NKp44, cells were nucleofected with a complex of Cas9 and sgRNA NCR2 (CRISPOR) at a ratio of 4:1. For the invalidation of perforin and LFA-1, cells were nucleofected with Cas9 and crRNA-tracRNA-ATTO550 (IDT) at a 1:1 ratio. For LFA-1 and perforin invalidation, 2 sgRNA were used by nucleofection.

### Complementation of KHYG-1 NKp44 KO with NKp44 transcripts

Lentiviral particles containing containing TRIP-DU3 encoding transcript for NKp44-1 (NCR2-201, CCDS CCDS56429; [ENST00000373083.8](https://www.ensembl.org/Homo_sapiens/Transcript/Summary?db=core;g=ENSG00000096264;r=6:41335608-41350889;t=ENST00000373083)), NKp44-2 (NCR2-202, CCDS CCDS56428; [ENST00000373086.3](https://asia.ensembl.org/Homo_sapiens/Transcript/Summary?db=core;g=ENSG00000096264;r=6:41335608-41350889;t=ENST00000373086)) or NKp44-3 (NCR2-203, BCCDS CCDS4855; [ENST00000373089.10](https://asia.ensembl.org/Homo_sapiens/Transcript/Summary?db=core;g=ENSG00000096264;r=6:41335608-41350889;t=ENST00000373089)) were generated using HEK 293FT cells as described in [81]. KHYG-1 NKp44KO Clone 1A1 cells were transduced with lentiviral particules in complete RPMI supplemented with 8µg/mL of polybrene (Sigma H9268). Cells were then maintained at 37°C in an atmosphere containing 5% CO_2_ for 3 days before assessing the expression of NKp44 by flow cytometry.

### sgRNA sequences used

sgRNA sequences used in this study are provided in the supplementary table.

### TIDE analysis

Genomic DNA from mutant and WT control cells was extracted using the NucleoSpin 96 Tissue Core Kit, a 96-well kit for DNA from cells and tissues (Macherey-Nagel) or the DNeasy Blood and Tissue Kit (Qiagen). DNA sequences from mutant and WT HCT-116 cells targeting the individual sgRNAs were amplified by PCR with specific primers and then sequenced. Primers were designed on Primer3 or IDT websites and selected to amplify amplicons of 500-1000bp in length. Each primer was designed to be at least 200 bp away from the sgRNA targeting site. PCR was migrated in a 1.3% agarose gel and amplicons were cut and purified using a gel and PCR clean-up kit (Macherey-Nagel) for further DNA sequencing. The DNA sequences were compared to analyze the indels (i.e. insertion or deletion) on the mutant DNA sequences compared to the WT DNA sequences using the TIDE (Tracking of Indels by Decomposition) website (http://shinyapps.datacurators.nl/tide/).

### Gene expression analysis

Total RNA was isolated from cell lines using a RNeasy mini kit (Qiagen). Reverse transcription was performed using Superscript RT III (Invitrogen). Amplification was performed using specific Taqman probes (Applied Biosystems) for each targeted gene (*Gapdh* Hs02786624_g1; *Pdgfd* Hs00228671_m1; *Nid1* Hs00915875_m1; *Hladpb1* Hs03045105_m1) and the Universal PCR Master Mix (Fluidigm). Samples were analysed on qPCR instrument (Applied Biosystems). Data were normalized (2^−ΔΔ*Ct*^) to the housekeeping genes, *Gapdh*.

### ICAM-1 expression

WT and SNP (rs1801714 C>T & rs5498 A>G) ICAM-1 transcript 1 (ICAM1-201, [ENST00000264832.8](https://asia.ensembl.org/Homo_sapiens/Transcript/Summary?db=core;g=ENSG00000090339;r=19:10271093-10286615;t=ENST00000264832)) and transcript 2 (ICAM-1-202, [ENST00000423829.2](https://asia.ensembl.org/Homo_sapiens/Transcript/Summary?db=core;g=ENSG00000090339;r=19:10271093-10286615;t=ENST00000423829)) were synthetized by Eurofins genomics. Plasmids were digested with BAMHI/Xho (NEB #R0136S & #R0146S) and cloned into lentiviral vector TRIPDU3 that have been digested with BamHI/XhoI. Subsequently, lentiviral particles were generated using HEK 293FT cells as described in [81]. Transductions were performed in 96W plates and cells were then maintained at 37°C in an atmosphere containing 5% CO_2_ for 3 days before assessing the expression of ICAM-1 by flow cytometry.

### Chromium release assay

Target cells were loaded with Chromium-51 (^51^Cr) (Perkin Elmer) for 1 h at 37°C under an atmosphere containing 5% CO_2_. Cells were washed 3 times in complete RPMI and plated out in triplicate (3,000 cells per well) with KHYG-1 cells at an E:T cell ratio of 10:1 in 96-well plates with U-bottom (BD Falcon). 10 µg/mL of the indicated monoclonal antibodies were added (in triplicate) and the plates were incubated for 4 hours at 37°C. After incubation, 50 μL of culture supernatant was transferred to a LumaPlate (Perkin Elmer) coated with solid scintillator, which was then placed in a microplate scintillation counter (Microbeta, Perkin Elmer) to determine the ^51^Cr into the supernatant, which was correlated with the lysis of target cells. The following formula was used to calculate the percentage of specific lysis: specific lysis (%) = (experimental release – spontaneous release) / (maximum release − spontaneous release) ×100. The maximum release was determined by adding 2% Triton X-100 (Sigma-Aldrich, 93443-100ML) or 2% Tergitol (Sigma-Aldrich, 15S9-100ML) to the target cells, and spontaneous release was measured in the medium alone, without effector cells.

### IncuCyte analysis

30,000 tumor cells were cultured with KHYG-1 at E:T cell ratio of 10:1 as described above in triplicate for 4 hours at 37°C and maintained at 37°C under an atmosphere containing 5% CO_2_. Cells were then washed and cultured at a dilution of 1/6 in complete RPMI medium containing 2 µg/mL puromycin in 96-flat bottom plates to allow attachment of target cells. 72 hours later, the plates were slowly washed twice to remove non-adherent cells (including dead KHYG-1 and dead HCT-116). The adherent target cells were then analyzed using the IncuCyte SC5 Live-Cell Analysis System (Sartorius). Plates were scanned using a 10x objective using phase contrast channel. Data were analyzed by counting adherent cells and normalized to untreated target cells to determine 100% tumor cell survival and confluence. Conditions were performed in triplicate and 5 images from each triplicate were used for analysis (15 in total). When indicated, 3,000 target cells were plated in triplicate with an increasing dose of recombinant human IFN-γ (Biolegend) in a 96-flat bottom plate for a final volume of 200 µL per well. Plates were scanned every 12 hours for 72 hours.

### Mice

C57BL/6J mice were purchased from JanvierLabs. *Ncr1^Cre^xR26^DTA^*and *Ncr1^CRE^xIfng^flox^* mice were bred in-house. Rosa26-DTA mice were purchased from Jackson laboratory lab (JAX:009669). *Ifng^flox^* mice were obtained from Alain Tedgui. Female mice were used at 8 weeks of age. All mice were bred in specific pathogen–free conditions at the Centre d’Immunologie de Marseille-Luminy and the Centre d’Immunophénomique (CIPHE). Experiments were conducted in accordance with institutional guidelines for the care and use of animals. All mouse experiments were performed in accordance with the rules of the CIML ethics committee and were approved by the *Ministère de l’Enseignement Supérieur, de la Recherche et de l’Innovation* – France (APAFIS#30510).

### B16F10 *in vivo* experiments

B16F10 cells were injected intravenously (i.v.) into the tail vein of C57BL/6 mice using a 30G syringe (0.5 × 10^5^ cells per mouse). The growth of lung tumor metastases was examined 10 days after injection. The mice were euthanized by cervical dislocation. The chest of the mice was examined postmortem and the lungs were harvested after perfusion with PBS 1X. Tumors were counted using Fiji software. For NK cell depletion, recipient mice were injected i.v. with 50 μg of anti-NK1.1 monoclonal antibody (clone PK136, Bio X Cell) two days before B16F10 injection and then once a week. mIgG2a monoclonal antibody was administered as a control. For IFN-γ neutralization, 200 µg/mouse anti-IFN-γ (clone XMG1.2, Bio X Cell) was injected intraperitoneally (i.p.) into the recipient mice one day before B16F10 injection and then every 2-3 days.

### Flow cytometry

#### Immune cell phenotyping

Human cells were incubated with normal mouse serum, and mouse cells were incubated with Fc block (anti-mouse CD16/CD32, BD) to saturate the Fc receptors, and then with the corresponding antibody cocktail diluted in staining buffer: 1X PBS (Gibco), 0.2% BSA (Sigma), 0.02% sodium azide (Prolabo) and 2 mM EDTA (Invitrogen). For human and mouse cell surface staining, cells were incubated with the indicated antibodies (see Table x). Cells were fixed in Cell Fix solution (BD) according to the manufacturer’s instructions. Data were acquired using an LSRFortessaX20 flow cytometer or a FACS Canto flow cytometer. The obtained FCS3.0 files were exported from BD FACSDiva software and imported into FlowJo v.10.5.2 (BD Biosciences).

#### Intracellular cytokine assessment

The splenocytes were cultured in 200 µL RPMI 1640 medium (Gibco) supplemented with 10% heat-inactivated fetal bovine serum (FBS, Gibco), 2 mM L-glutamine (Gibco), 1% non-essential amino acids (Gibco), 1 mM sodium pyruvate (Gibco) and 1% Hepes 1M (Gibco) and 1% penicillin-streptomycin (10 000 U/ml, Gibco) with phorbol 12-myristate 13-acetate (PMA, 125 ng/mL final; Sigma, P8139) and ionomycin (IONO, 1 µg/mL final; Sigma, I0634) and GolgiStop (BD, 1/1500) to increase intracellular protein transport. and were kept for 4 hours at 37°C under an atmosphere of 5% CO_2_. Control conditions were obtained without PMA and ionomycin. The cells were then washed twice in staining buffer and incubated with the antibodies anti-CD3-Pacific Blue (BD), anti-CD8, anti-NK1.1 and anti-NKp46 (clone 29A1.4 Biolegend) for 45 min at 4°C. The cells were washed twice, fixed and permeabilized with Cytofix/Cytoperm (BD). The cells were washed twice in Perm/Wash (BD) and stained by incubation with anti-IFN-γ (BD) antibody for 30 min at 4°C. Cells were washed twice and data were acquired using the LSR2UV flow cytometer.

#### Cytochrome C release

150,000 KHYG-1 cells were incubated with 30,000 indicated target cells in a 5:1 ratio in triplicate at a final volume of 200 µL per well in 96-well plates with U-bottom (BD Falcon) for 4 hours at 37°C and maintained at 37°C under an atmosphere containing 5% CO_2_. Cells were then washed 1 time with complete medium and cultured in complete medium containing 2 µg/mL puromycin (Gibco) at a final volume of 200 µL per well in flat-bottom plates and maintained at 37°C under an atmosphere of 5% CO_2_ for 24 hours. Cells were washed twice in staining buffer and incubated with Live/Dead Fixable Near IR Dead Cell Stain Kit (Invitrogen) and anti-human CD45-BV786 in staining buffer for 20 min at 4°C. Cells were then washed twice and fixed/permeabilized (Ebioscience fix/perm buffer) for 20 min. The cells were then washed with permeabilization buffer (PBS1X 10% BSA 0.5% Tween) and incubated with anti-cytochrome C in permeabilization buffer overnight at 4°C. The cells were analyzed directly with the LSRFortessaX20 flow cytometer.

## SUPPLEMENTARY MATERIALS

**Figure S1:**
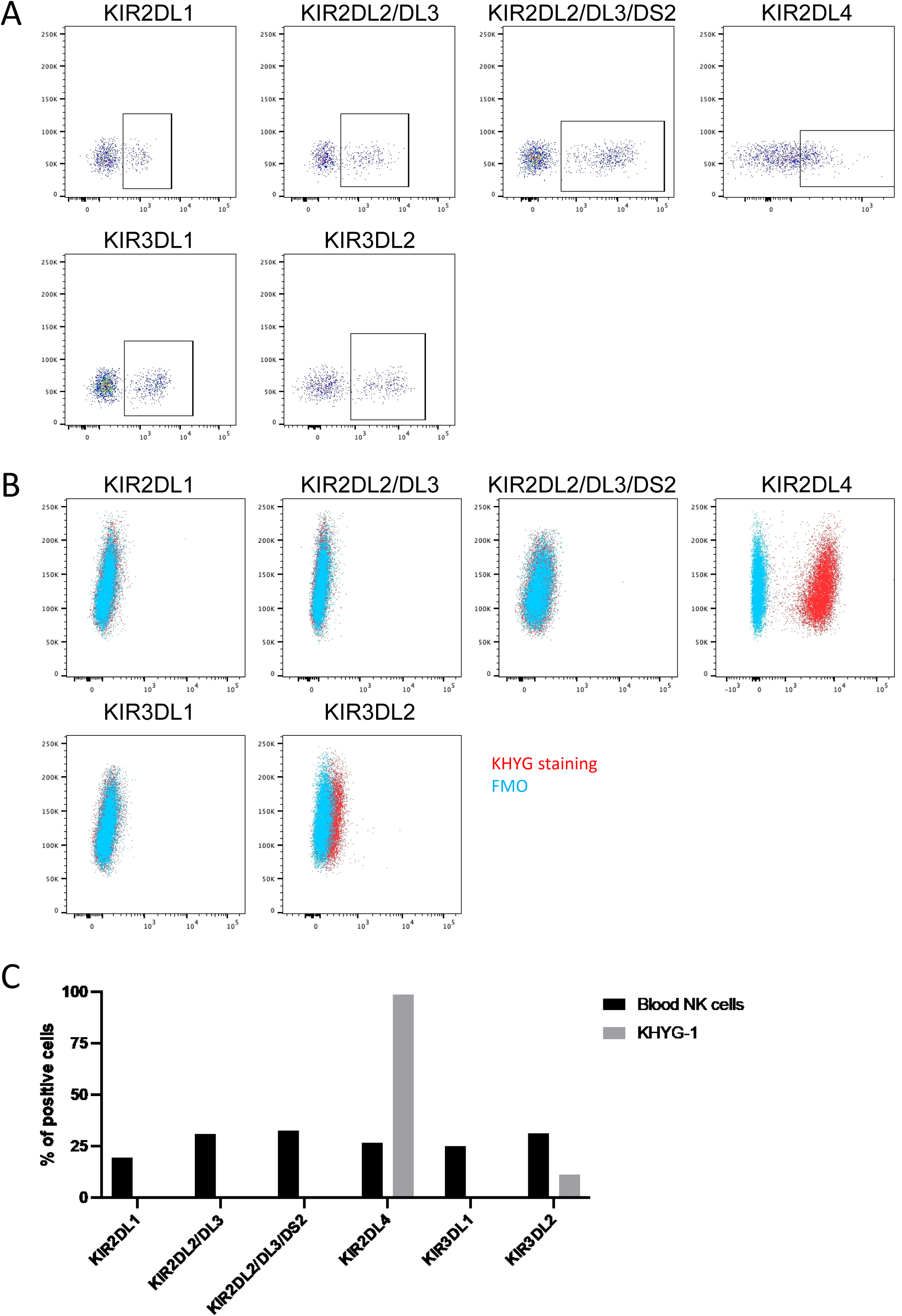
HCT-116 tumor cells do not express inhibitory KIR. FACS profiles of indicated KIR at the cell surface of NK cells from peripheral blood mononuclear cells (PBMC) (A) and HCT-116 cells (B). In (C) % of positive cells are shown.

**Figure S2:**
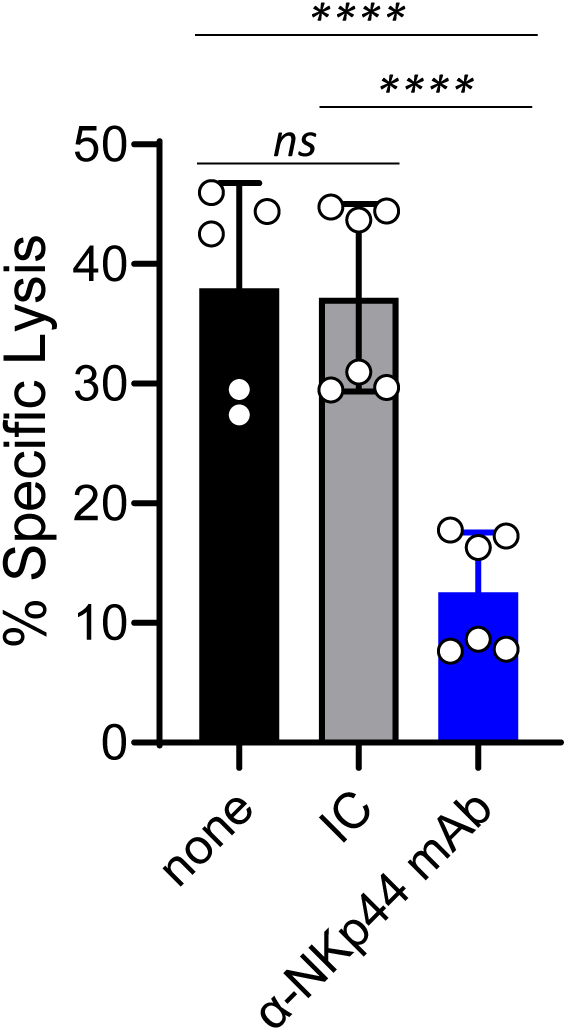
HCT-116 tumor cells are susceptible to NK cell-mediated lysis by NKp44. HCT-116 cells were co-cultured with KHYG-1 cells in the presence or absence of an anti-NKp44 mAb or isotype control. The extent of specific lysis was determined in a standard 4-hour chromium release assay. Data result from the pool of 2 independent experiments and are representative of more than 10 experiments. One-way anova with multiple comparison. Adjusted P value ****=0.0001

**Figure S3:**
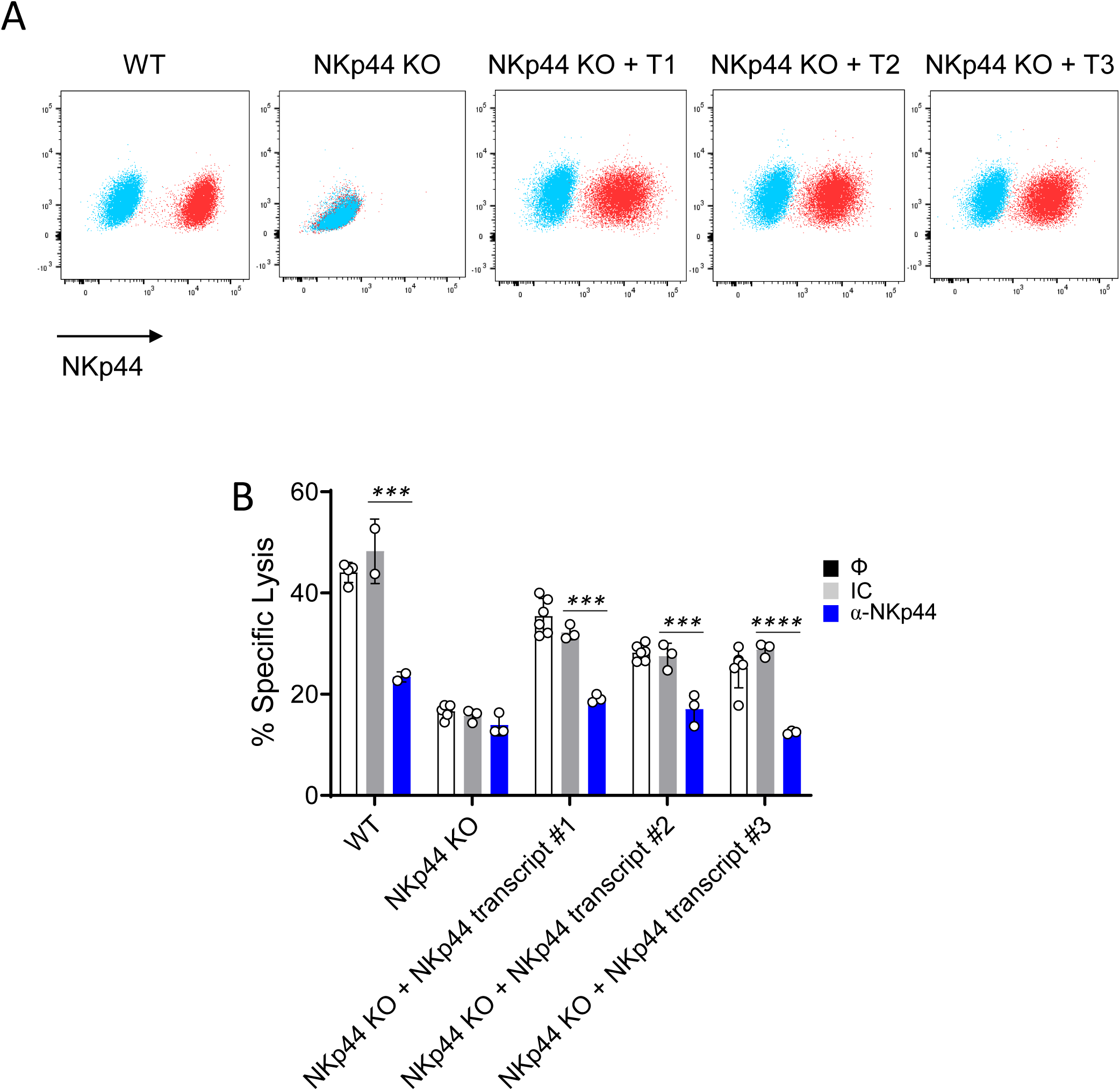
Expression of NKp44 encoding transcripts in NKp44 KO KHYG-1 cells restore NK cell-mediated lysis of HCT-116 tumor cells by NKp44. (A) FACS profile of NKp44 cell surface expression on KHYG-1 cells, NKp44 KO KHYG-1 cells and NKp44 KO KHYG-1 cells transduced with a lentiviral vector encoding the indicated NKp44 transcript. (B) HCT-116 cells were co-cultured with indicated KHYG-1 cells. An anti-NKp44 mAb or isotype control was added as indicated. The extent of specific lysis was determined in a standard 4-hour chromium release assay. Data are representative of 3 independent experiments. One-Way anova with SidaK’s multiple comparison tests. ***p<0.005; ****p<0.0001.

**Figure S4:**
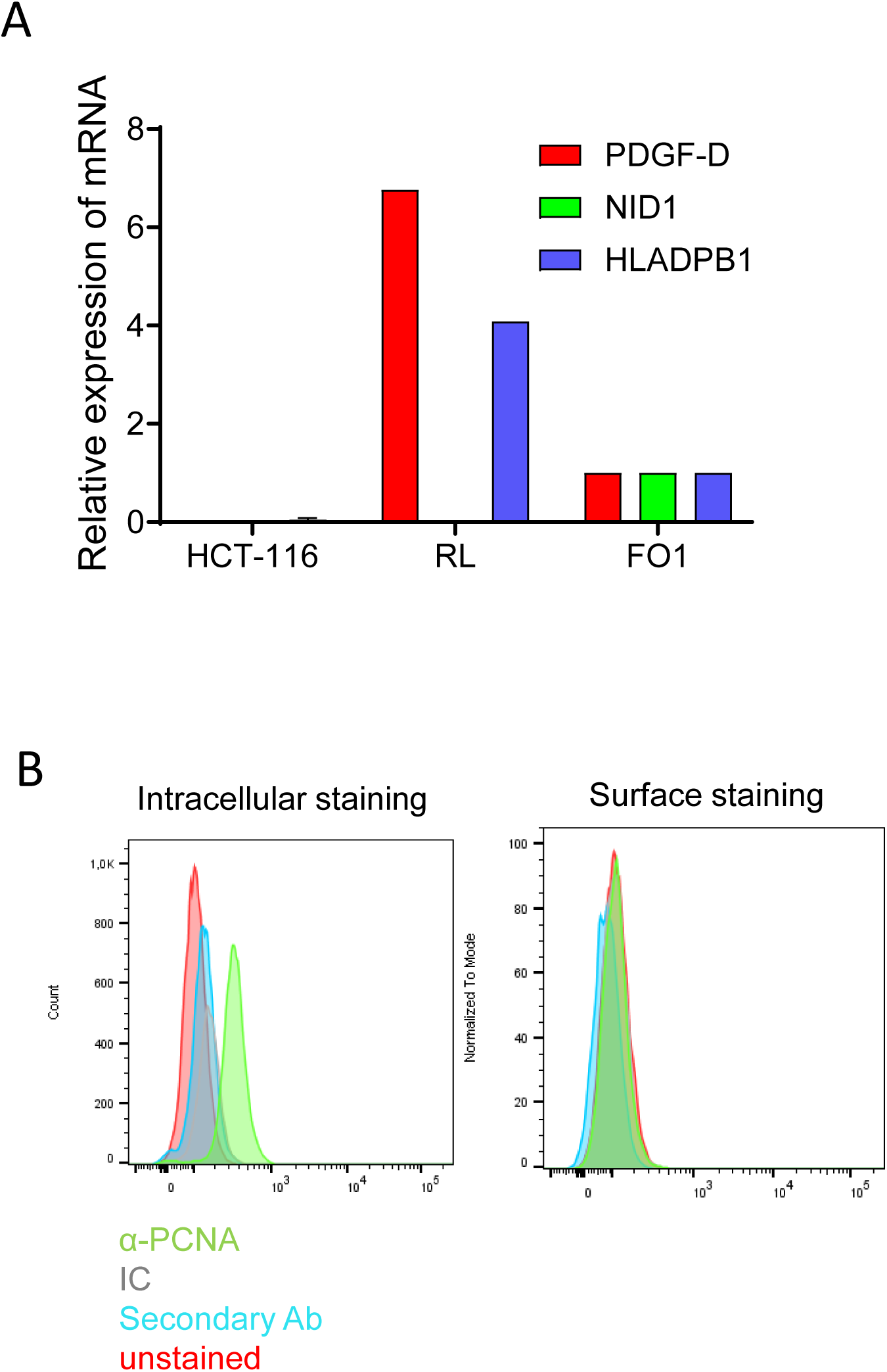
HCT-116 cells do not express the described NKp44 ligands. (A) Relative mRNA expression of indicated NKp44 ligand in HCT-116, RL and FO1 cells. (B) FACS profiles of cell surface and intracellular expression of PCNA. Data are representative of 2 independent experiments.

**Figure S5:**
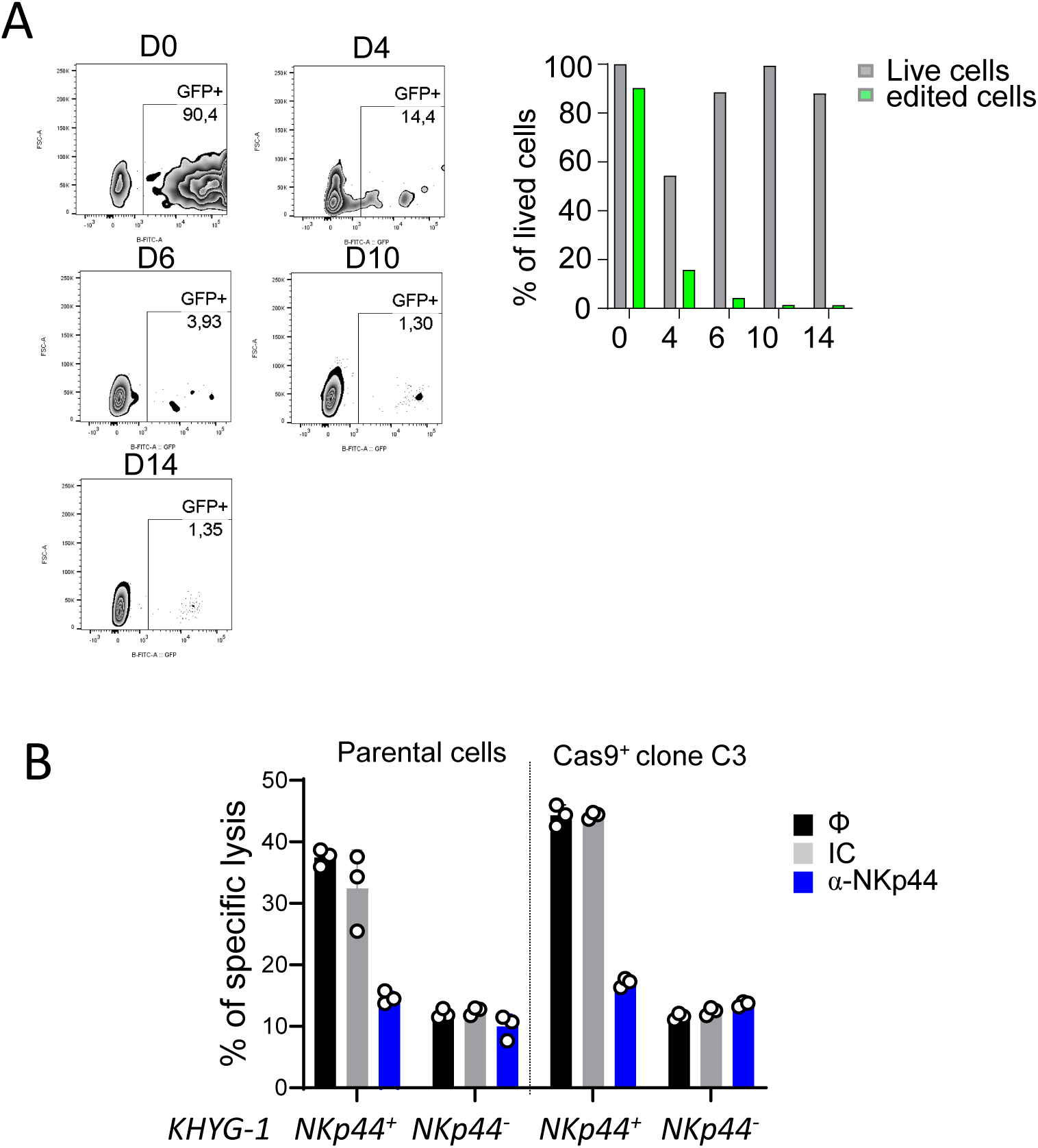
Selection of HCT-116 Cas9^+^ clones that efficiently edit the genome. HCT-116 cells were transduced with an inducible Cas9 encoding vector and cloned. Several clones were expanded and a portion of them were transduced with a lentiviral vector encoding sgRNA targeting the survival gene RPL6 and expressing GFP. Gene editing was monitored by tracking the decrease in the number of GFP^+^ cells over time. In (A), data show clone C3. (B) HCT-116 Cas9^+^ clone C3 was cocultured with indicated WT or NKp44 deficient KHYG-1 cells for 4 hours and specific lysis was determined in a standard chromium release assay.

**Figure S6:**
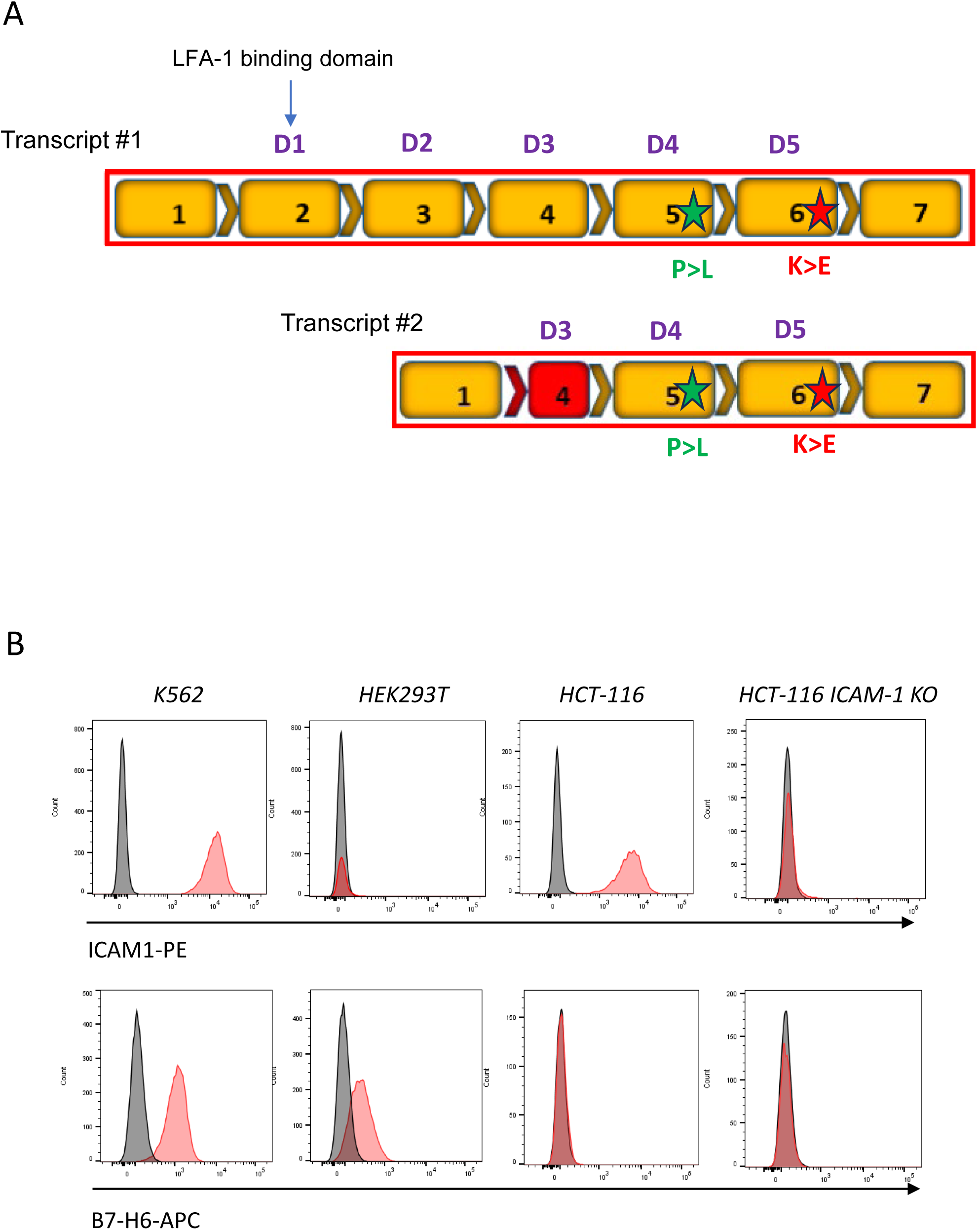
Cell surface expression of ICAM-1 and B7-H6 on tumor cells. (A) Schematic representation of ICAM-1 transcripts #1 and #2 with exons in orange. The domains (D1 to D5) are indicated. SNP are indicated by a star. (B) FACS profile of ICAM-1 and B7-H6 cell surface expression on HCT-116, ICAM-1-deficient HCT-116, K562 and HEK 293T cells. Data are representative of at least 2 independent experiments.

**Figure S7:**
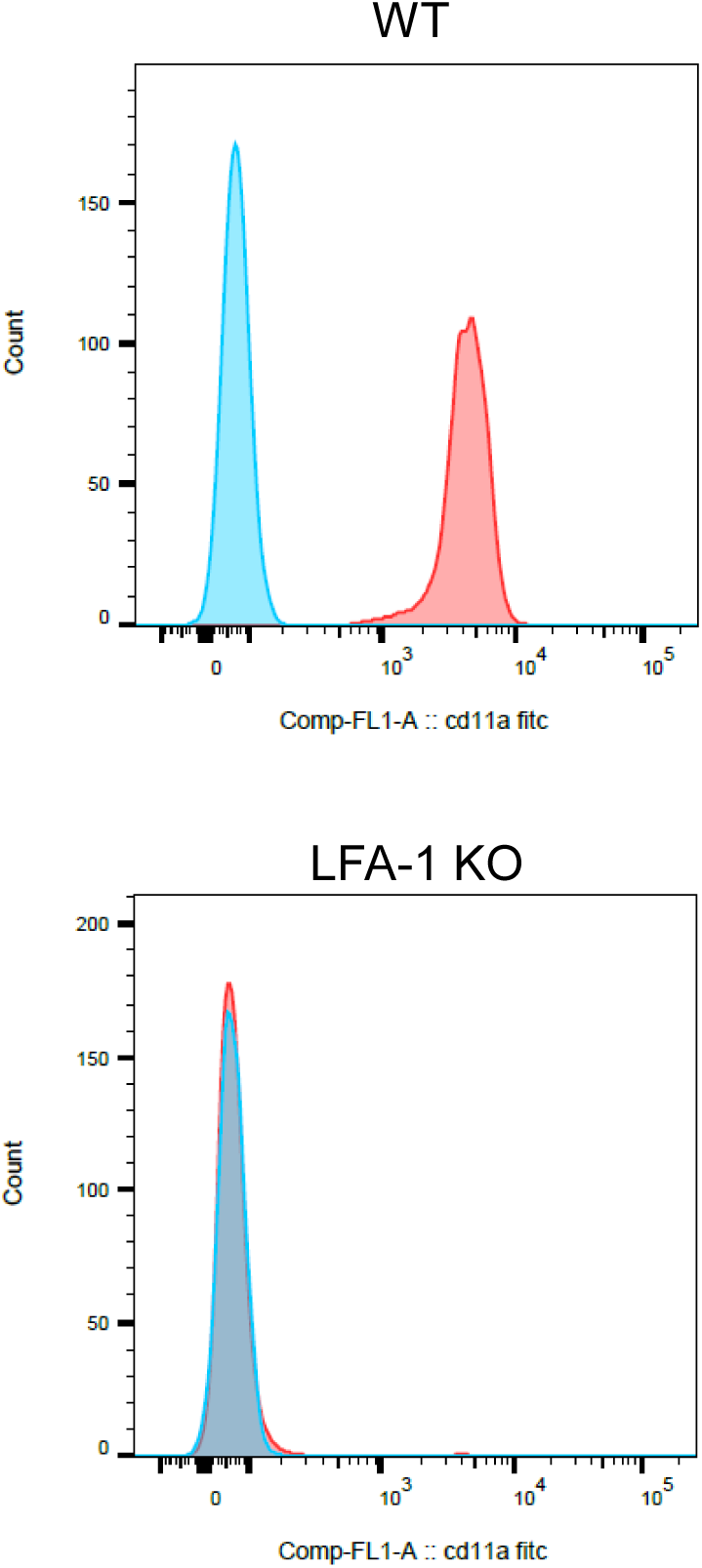
Cell surface expression of LFA-1 in KHYG-1 and LFA-1 KO KHYG-1 cells. (A) FACS profile of LFA-1 (CD11a) cell surface expression on KHYG-1 and LFA-1 KO KHYG-1 cells. Data are representative of at least 3 independent experiments.

**Figure S8:**
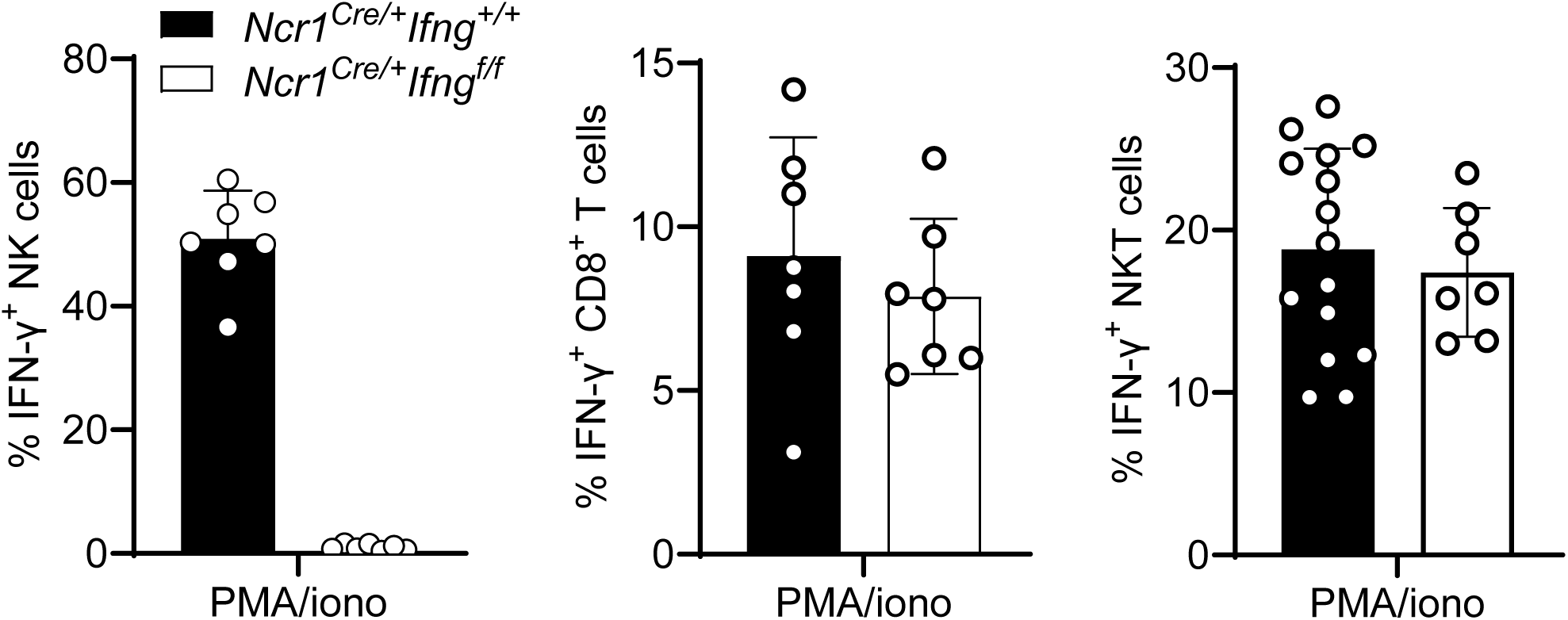
IFN-γ production by NK cells, CD8^+^ T cells and NKT cells from *Ncr1^iCre/+^Ifng^fl/fl^* after in vitro stimulation. Splenocytes from *Ncr1^iCre/+^Ifng^fl/fl^*and control *Ncr1^iCre/+^Ifng^+/+^* mice were stimulated with PMA/inomycin. Frequencies of IFN-γ producing NK cells, CD8^+^ T cells and NKT cells after stimulation are shown. Data result from the pool of 3 independent experiments.

